# Maintenance and transformation of representational formats during working memory prioritization

**DOI:** 10.1101/2023.02.08.527513

**Authors:** Daniel Pacheco Estefan, Marie Christin Fellner, Lukas Kunz, Hui Zhang, Peter Reinacher, Charlotte Roy, Armin Brandt, Andreas Schulze-Bonhage, Linglin Yang, Shuang Wang, Jing Liu, Gui Xue, Nikolai Axmacher

## Abstract

Visual working memory depends on both material-specific brain areas in the ventral visual stream (VVS) that support the maintenance of stimulus representations and on regions in prefrontal cortex (PFC) that control these representations. Recent studies identified stimulus-specific working memory contents via representational similarity analysis (RSA) and analyzed their representational format using deep neural networks (DNNs) as models of the multi-layered hierarchy of information processing. How executive control prioritizes relevant working memory contents and whether this affects their representational formats remains an open question, however. Here, we addressed this issue using a multi-item working memory task involving a retro-cue that prompted participants to maintain one particular item. We exploited the excellent spatiotemporal resolution of intracranial EEG (iEEG) recordings in epilepsy patients and analyzed activity at electrodes in VVS (n=28 patients) and PFC (n=16 patients). During encoding, we identified category-specific information in both VVS and PFC. During maintenance, this information re-occurred in VVS but not in the PFC – suggesting a transformation of PFC representations from encoding to maintenance which putatively reflects the prioritization process. We thus applied RSA in combination with different DNN architectures to investigate the representational format of prioritized working memory contents. Representations during the maintenance period matched representations in deep layers of recurrent but not feedforward DNNs, in both VVS and PFC. While recurrent DNN representations matched PFC representations in the beta band following the retro-cue, they corresponded to VVS representations in a lower theta-alpha frequency range (3-14Hz) towards the end of the maintenance period. Findings could be replicated in recurrent DNNs with two different architectures and using two different training sets. Together, these results demonstrate that VWM relies on representational transformations in VVS and PFC that give rise to distinct coding schemes of prioritized contents.

## Introduction

Visual working memory (VWM) refers to the ability to store visual information for a short period of time and to flexibly manipulate this information according to task demands. One essential aspect of VWM memory is prioritization, i.e., the ability to selectively allocate attention to particular features or items depending on behavioral or cognitive requirements. Influential theories have proposed that WM prioritization entails the transformation of maintained representations from a purely mnemonic to a task-optimized state (Chatham et al., 2014; Myers et al., 2017). On a neurophysiological level, these accounts predict that working memory prioritization involves a task-dependent transformation of representational patterns in executive control areas which can be disentangled from a mnemonic coding scheme that maintains perceptual stimulus features in sensory brain regions (Liebe et al., 2012; Miller et al., 2018; Myers et al., 2017; Stokes et al., 2013). Here we set out to test this prediction. We analyzed the representational format of VWM stimuli using electrophysiological recordings in human epilepsy patients implanted with electrodes in ventral visual stream (VVS) and/or prefrontal cortex (PFC).

There is now abundant evidence on the neural correlates of VWM control processes in humans (Lepsien et al., 2011; Lepsien & Nobre, 2006; Nee & Jonides, 2008, 2009). Early studies focused on prioritization of to-be-encoded items, using paradigms in which participants were asked to selectively attend to particular items before these items were shown (e.g., Griffin & Nobre, 2003; Posner, 1980; Schmidt et al., 2002; Vogel & Machizawa, 2004). More recent investigations often employed retrospective cueing paradigms, in which prioritization is applied to information after its encoding into VWM (Ester et al., 2018; Higo et al., 2011; Nee & Jonides, 2009; Nelissen et al., 2013; Sprague et al., 2016). These studies revealed that prefrontal and parietal regions which underlie the allocation of attention during perception are also engaged in the prioritization of items in VWM (Buschman & Kastner, 2015; Buschman & Miller, 2007; Chatham et al., 2014; D’Esposito, 2007; Lepsien et al., 2011; Lepsien & Nobre, 2006; Nee & Jonides, 2008; Nobre et al., 2004). Notably, a recent meta-analysis observed selective responses to retro-cues but not to cues that allocate attention prior to item encoding in VWM in various prefrontal areas (Myers et al., 2017; Wallis et al., 2015). This literature suggests that prioritization affects VWM representations in the PFC, yet this prediction has not been tested experimentally in humans.

The critical role of the PFC in WM prioritization is commonly believed to depend on dynamic recurrent computations. A classical model of WM suggests that persistent activity depends on reverberatory excitation within a local recurrent neural network (Barak & Tsodyks, 2014; X. J. Wang, 2001). Computational studies have shown that recurrence is crucial for the selection and integration of task-relevant features in the PFC (Mante et al., 2013), the integration of working memory and planning (Ehrlich & Murray, 2022), the flexibility of WM and the avoidance of interference in the presence of competing representations (Bouchacourt & Buschman, 2019), and – most importantly for our study – WM prioritization (Wan et al., 2022). Recurrent computations might be particularly relevant for selective attention to specific features or items in WM because they enable the stabilization of reverberating activity in attractor states that modulate the excitability of assemblies which represent prioritized contents (Barak & Tsodyks, 2014; Compte et al., 2000; X. J. Wang, 2001). In addition to their theorized role in PFC prioritization, recurrent computations have been proposed to be critical for information processing in the VVS during visual perception (Kar & DiCarlo, 2021; Kietzmann et al., 2019), and for offline ‘generation’ of stimuli during visual imagery (Breedlove et al., 2020). However, a specific role of recurrency in the VVS for VWM maintenance has not been previously investigated.

In addition (and possibly related) to the relevance of recurrent computations, theories have emphasized the important role of brain oscillations for VWM, in particular for the prioritization process. Oscillatory activity in the gamma frequency range (50-120Hz) is thought to convey bottom-up information during VWM encoding, while oscillations at beta frequency (20-35Hz) are supposed to provide top-down control over VWM contents (Engel & Fries, 2010; Kuzovkin et al., 2018; Lundqvist et al., 2018; Miller et al., 2018; Vezoli et al., 2021). The significance of these oscillatory patterns has been validated experimentally in a series of studies in macaques (Buschman & Miller, 2007; Lundqvist et al., 2018; Miller et al., 2018). Furthermore, a recent study in humans confirmed the crucial role of gamma-band activity (30-75Hz) for conveying bottom-up information from lower-level visual areas to regions processing higher-level information (Kuzovkin et al., 2018). In addition to their role in top-down control over WM content, several studies have now associated beta oscillations with the reactivation of stimulus-specific activity during the VWM prioritization process. Content-specific beta activity has been shown to carry information about internalized task rules (Buschman et al., 2012), stimulus categories (Antzoulatos & Miller, 2014, 2016; Stanley et al., 2018), scalar magnitudes (Spitzer et al., 2010, 2014) and perceptual decisions (Wimmer et al., 2016); for review, see (Spitzer & Haegens, 2017). These studies highlight the role of beta oscillations in encoding task-relevant stimulus properties.

In humans, intracranial EEG (iEEG) recordings in epilepsy patients have been used to investigate the neurophysiological patterns underlying content-specific memory representations. This research has employed multivariate analysis techniques, such as pattern classification and representational similarity analysis (RSA), to identify representations of specific stimuli (Kriegeskorte et al., 2008; Kriegeskorte & Diedrichsen, 2019). Studies have demonstrated that frequency-specific representations in the gamma, beta and theta (3-8Hz) frequency bands contain item- and category-specific information, playing a crucial role in episodic memory retrieval (Pacheco Estefan et al., 2019, 2021). In addition to identifying the relevant oscillatory frequencies that carry representational content during visual perception and episodic memory, recent iEEG studies have investigated the ‘formats’ of VWM representations. This research employed deep neural networks (DNNs) to investigate how different aspects of natural images are represented in the brain during mnemonic processing. These studies assume that mnemonic representations require specialized circuits for processing distinct aspects (or formats) of natural images, from low-level sensory features to higher-level contents and conceptual/semantic information (Axmacher, 2020; Heinen et al., 2023; Kwak & Curtis, 2022; W. Tang et al., 2023; Wu & Fuentemilla, 2023; Xue, 2022). Indeed, several studies assessed the different representational formats during VWM encoding and maintenance and demonstrated substantial transformations of VWM representations into a format that aligns with late layers of a convolutional DNN (Liu et al., 2020, 2021). While these results and methodological advancements have provided valuable insights into the format of VWM representations, no study so far has investigated the representational transformations that accompany VWM prioritization in humans. Thus, whether the prioritization of VWM representations involves a change in the representational format of the stored content and distinct coding schemes of attended (i.e., task-relevant) items is currently unknown.

Here, we leveraged the heuristic potential of DNNs as models of visual representation, the flexibility of RSA, and the high spatiotemporal resolution of iEEG to investigate this topic. We analyzed electrophysiological activity from VVS and PFC while patients performed a multi-item VWM paradigm involving a retro-cue. Participants encoded a sequence of three images and were then prompted by a retro-cue to maintain either one of these items or all items (Figure 1A; Methods). The objects belonged to six categories, each containing ten exemplars (60 images in total; Figure 1B). With the exception of the behavioral data (Figure 1D), we only analyzed activity during the single item condition in this study, given our focus on the prioritization process. This experimental design allowed us to evaluate how information about specific contents is represented in the brain during initial encoding and how it is transformed due to the retro-cue, both in terms of representational formats and regarding the frequencies of brain oscillations in VVS (438 electrodes) and PFC (146 electrodes; Figure 1C). We hypothesized that frequency-specific representations would reflect bottom-up storage and top-down information transfer, respectively, with a particular role of gamma and beta oscillations (Engel & Fries, 2010; Kuzovkin et al., 2018; Lundqvist et al., 2018; Miller et al., 2018; Vezoli et al., 2021). Specifically, we predicted that oscillatory PFC activity in the beta frequency range may reflect representational transformations due to top-down control following the retro-cue (Miller et al., 2018). In addition, we expected recurrent convolutional architectures to better explain representations than feedforward DNNs during VWM maintenance (Bouchacourt & Buschman, 2019; Kar et al., 2019; Kar & DiCarlo, 2021; Kietzmann et al., 2019; van Bergen & Kriegeskorte, 2020; X.-J. Wang, 2001).

**Figure 1:**
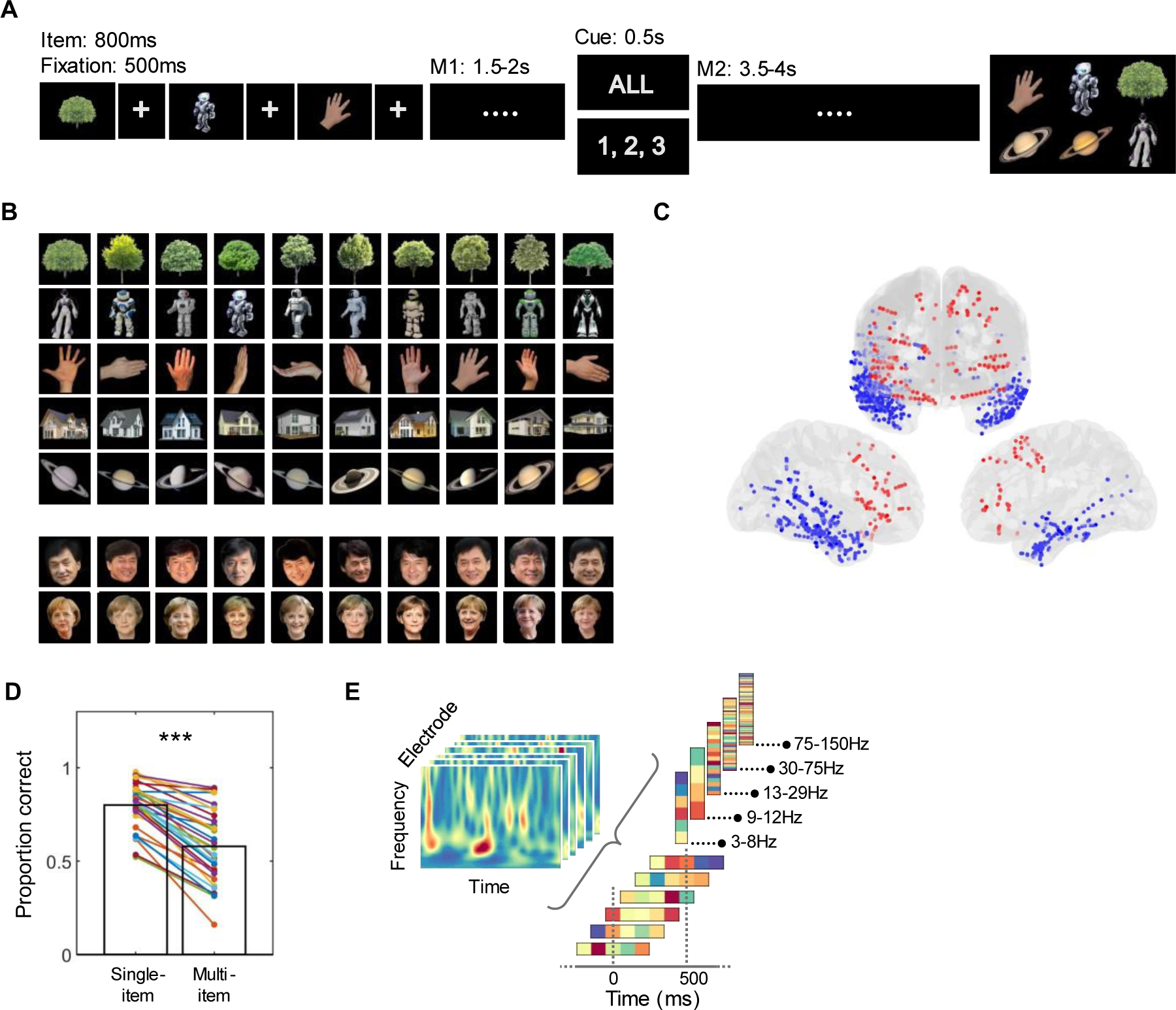
Experimental procedure, electrode coverage, and behavioral results. (A) Participants encoded a sequence of 3 images of natural objects and were asked to remember this content during two subsequent maintenance periods that were separated by a retro-cue (M1 – retro-cue – M2). The cue prompted participants to either selectively maintain items at particular list positions (single-item trials, “1, 2, 3” in the figure) or to maintain all items in their order of presentation (multi-item trials, “All” in the figure). During the probe, six items were presented, which included all encoded items, two novel exemplars from previously presented categories, and one item from a novel category. (B) All images presented in the experiment (60 in total). Objects pertained to six categories with ten exemplars each. Two different versions of the experiment were created for different patient populations, including a different famous face (bottom). (C) Electrode implantation included 438 electrodes in the ventral visual stream (VVS, N = 28 participants; blue) and 146 electrodes in the prefrontal cortex (PFC, N = 16 participants; red). (D) Behavioral performance was significantly higher for single as compared to multi-item trials. (E) Representational patterns included in the RSA analyses were computed in windows of 500ms, incrementing in 100ms steps, using Spearman’s correlation (rho) across electrodes, time points (5 time points in each 500ms window), and frequencies. Analyses were performed in individual frequencies in the 3-150Hz range in the model-based RSA analyses, and within different frequency bands (theta, alpha, beta, low gamma, high gamma) in the contrast-based pattern similarity analyses (Methods). ***: p < 0.001

## Results

### Behavioral results

Successful prioritization in the single-item condition should result in better performance than in the multi-item trials. Indeed, we found that participants performed significantly better in single-item trials (proportion correct trials: 0.80 ± 0.12) than in the multi-item condition (0.58 ± 0.19; t(31) = 11.23; p = 1.84e-12; Figure 1D). This suggests that participants followed instructions and benefited from prioritizing task-relevant representations in the single-item trials.

### Maintenance and transformation of category-specific representations

We investigated the electrophysiological patterns supporting the representation of category-specific information in VVS and PFC. As a first approach, we assessed the presence of categorical representations, employing RSA and a simple model of category information (Figure 2A). We constructed an item-by-item representational similarity matrix (RSM) reflecting the hypothesis that items of the same category would elicit more similar patterns of brain activity compared to items of different categories (Figure 2A). We correlated this model RSM with temporally resolved neural RSMs (windows of 500ms, overlapping by 400ms). Representational patterns included power values across electrodes (16.85 ± 8.92 electrodes in VVS, 9.73 ± 11.1 in PFC; Mean ± STD) and time points (5 time points of 100ms in each time window; Figure 1E, Methods) and were analyzed separately in each of 52 different frequencies between 3 and 150 Hz. To determine the similarity between feature vectors, we used Spearman’s Rho (Liu et al., 2020; Pacheco Estefan et al., 2021).

**Figure 2:**
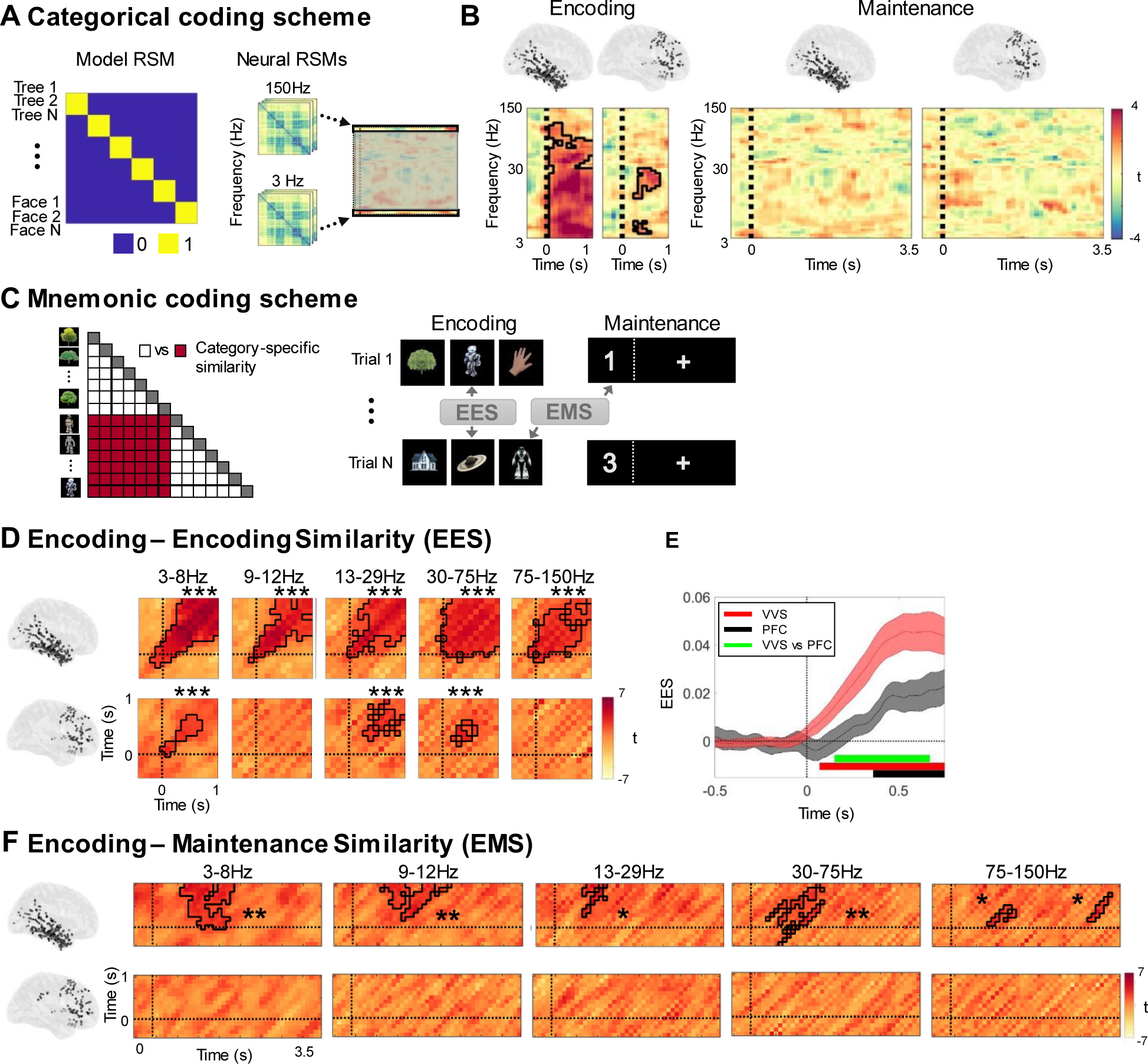
Encoding and maintenance of category-specific representations. (A) Categorical coding scheme. Representational similarity matrix (RSM) reflecting the hypothesis that category information structures the representational geometry of stimuli (left) were correlated with a time-series of neural RSMs (right) at each individual frequency. (B) Fit of the category model in the VVS and the PFC during the encoding (left) and maintenance (right) period. (C) Mnemonic coding scheme. Left: Category-specific similarity was computed by contrasting correlations between (different) items of the same category vs items of different categories. Right: Similarity was calculated between the encoding periods of different trials (encoding-encoding similarity, EES) or between the encoding and the maintenance periods of different trials (encoding-maintenance similarity, EMS). (D) EES analysis: category contrasts for five frequency bands during encoding in VVS (top) and PFC (bottom). Clusters with significant differences between same and different categories surviving correction for multiple comparisons using cluster-based permutation statistics are outlined in black. Zero indicates the onset of image presentation on both time axes. (E) Category specificity analysis at higher temporal resolution showing different latencies of effects in VVS (red) and PFC (black). Each line shows the time course of within minus between category correlations in each region. Horizontal bars indicate time-periods when EES values are significantly different from zero in each region, and significantly different between the two regions (green). (F) Encoding-Maintenance Similarity was computed by correlating patterns of activity at different encoding time windows (vertical axis) with maintenance activity after presentation of the retro-cue (horizontal axis), for the VVS (top) and the PFC (bottom). Zero indicates image onset during encoding and retro-cue onset during maintenance, respectively. ***: p < 0.001; * : p < 0.01; * : p < 0.05.

Our simple category model revealed a marked presence of categorical information during encoding in the VVS. This was observed in a significant frequency cluster ranging from 3-120Hz that started immediately after stimulus presentation and lasted for the whole encoding period (0.8s; p = 0.001). In the PFC, we observed two clusters of significant fit in the beta (17-28Hz; 200-800ms; p = 0.001) and the theta frequency range (3-7Hz; 200-600ms; p = 0.044; Figure 2B, left). During maintenance, we did not observe any significant fits between model and neural RSMs in either VVS or PFC (VVS: all p > 0.51; PFC: all p > 0.105; Figure 2B, right).

The absence of fit of the category model during maintenance might be attributed to a weakening of the representations during the maintenance period – e.g., due to a decrease in signal to noise ratio – or to a rapid transformation of activity patterns during encoding (Liu et al., 2021). To evaluate whether transformed activity patterns from encoding reoccur during the maintenance period, we performed a category-specific pattern similarity analysis (Methods; Liu et al., 2020). This analysis involved contrasting correlations of items belonging to the same category with correlations of items from different categories (Figure 2C, left), both during encoding (encoding-encoding similarity; EES) and between encoding and maintenance (encoding-maintenance similarity, EMS; Figure 2C, Right). Notably, while the category model can track the presence of categorical representations at the level of the representational geometry of our stimuli set, the EES and EMS analyses test for re-occurrence of category-related neural activity patterns from different encoding periods. This analysis was conducted in 5 conventional frequency bands (theta, 3-8Hz; alpha, 9-12Hz; beta, 13-29Hz; low gamma, 30-75Hz; high gamma, 75-150Hz), with electrodes, time points (including both matching and non-matching time points; see Methods), and frequencies in each band as features.

We first analyzed the timing and temporal stability of representations during encoding, using EES. Consistent with the results observed in the category model analysis, the EES analysis revealed prominent category-specific information during the encoding phase in both VVS and PFC. In the VVS, this was observed in all frequency bands (all p_corr_ < 0.005; Figure 2D, top). In the PFC, category-specific information was found in the theta, beta and low-gamma frequency bands (all p_corr_ < 0.01; Figure 2D, bottom). To examine the relative timing of the effects in greater detail, we increased the temporal resolution by shortening the sliding windows to steps of 10ms and including all frequencies in the 3-150Hz range as RSA features. This analysis demonstrated that categorical information reached the PFC 360ms after stimulus presentation, i.e., 290ms later than the VVS (Figure 2E). Indeed, a direct comparison of latencies showed significantly higher EES in VVS than PFC starting 150ms after stimulus onset (p_corr_ = 0.007). Furthermore, in the majority of frequency bands category-specific representations were most pronounced at matching time points across trials (diagonal values in Figure 2D) and did not generalize to other time periods, in line with theories on dynamic coding (Liu et al., 2020; Stokes et al., 2013).

Next, we analyzed Encoding-Maintenance Similarity (EMS) to investigate whether category-specific representations established during encoding reoccurred during the maintenance phase. We observed significant reoccurrence of category-specific information in all frequency bands in the VVS (all p_corr_ < 0.025). These effects were transient and most pronounced within the first 2 seconds after the retro-cue (Figure 2F, top). In contrast, we did not observe reoccurrence of category-specific information in the PFC in any frequency band (all p_corr_ > 0.19; Figure 2F, bottom).

Together, these results show the formation of category-specific representations in both VVS and (later) in PFC during encoding, but reoccurrence of encoding patterns during maintenance in the VVS only.

### Representational formats of category-specific representations

Our findings presented thus far indicate maintenance of category-specific representations from encoding in the VVS that was not observed in the PFC. The absence of an effect in the PFC may be attributed to a transformation of VWM representations driven by the prioritization process. Indeed, recent behavioral (Kerren et al., 2022), neuroimaging (Kwak & Curtis, 2022) and iEEG studies (Liu et al., 2020, 2021) established a crucial role of transformed representational formats, particularly abstract representational formats devoid of specific sensory information, in VWM maintenance. Based on these insights, we hypothesized that the PFC may contain categorical information in an abstract representational format after presentation of the retro-cue.

To evaluate this hypothesis, we employed different deep neural network (DNN) architectures. First, we used the feedforward DNN ‘AlexNet’ (Krizhevsky et al., 2012) that has been extensively employed to characterize neural representations of natural images during perceptual and mnemonic processes (Baek et al., 2021; Bao et al., 2020; Cadieu et al., 2014; Cichy et al., 2016; Khaligh-Razavi & Kriegeskorte, 2014; Kuzovkin et al., 2018; Lindsay, 2021; Liu et al., 2020, 2021; H. Tang et al., 2018; Vinken & Op de Beeck, 2021). Additionally, we applied two recurrent DNNs, the BL-NET and the corNET-RT. The BL-NET consists of seven convolutional layers which include lateral recurrent connections and has previously been applied to predict human behavior, specifically reaction times, in a perceptual task (Spoerer et al., 2020). The corNET-RT has a relatively shallow architecture compared to similarly performing networks for image classification and has been designed to model information processing dynamics in the primate VVS (Kubilius et al., 2018). Similar to the BL-NET, corNET-RT exhibits recurrent dynamics that propagate within (but not between) layers. All 3 DNNs represent stimuli in various representational formats, ranging from low-level visual features in superficial layers to conceptual and abstract features in deep layers. While AlexNet processes stimulus features in a single feedforward pass, the lateral recurrent connections of BL-NET and corNET-RT generate temporally evolving time-series of stimulus representations in each layer, thus capturing core properties of recurrent dynamics during WM processing in the PFC. The number of recurrent passes is fixed to 8 time-points in BL-NET, while the corNET-RT model exhibits layer-specific recurrent passes that range from 2 to 5 time points (see Methods). Following previous studies, and in order to ensure that the networks achieved stable representations of our images in each layer, we focused on the RSMs at the last time-point of each layer (Muttenthaler & Hebart, 2021).

We first characterized stimulus representations in different layers of AlexNet. We constructed RSMs from DNN representations by computing the similarities between all unit activations in each layer for all pairs of images (Liu et al., 2020, 2021; Figure 3A, top). For visualization, we projected the data into two-dimensional space using Multidimensional Scaling (Figure 3A, bottom). To evaluate representational changes throughout the DNN, we correlated the RSMs between different layers. RSMs were most similar among the convolutional layers 2-5 and among the fully connected layers 6-7, while the input layer 1 and the output layer 8 exhibited the most distinct representational patterns (Figure 3B). We computed the Category Cluster Index (CCI; McKee et al., 2014; Mehrer et al., 2020), defined as the difference in average distances of stimulus pairs from the same category vs. stimulus pairs from different categories (Figure 3C). CCI takes a value of 1 if clusters are exclusively built by stimuli from the same category and approaches 0 if the representational geometry shows no categorical organization. Using permutation statistics (i.e., label shuffling), we observed that CCI values were significantly higher than chance in all layers of the network (all p_corr_ = 0.008, Bonferroni-corrected for the 8 layers). Notably, we observed a four-fold increase in CCI scores from the first (CCI = 0.11) to the last layer (CCI = 0.46) of the AlexNet (Figure 3C). This effect was explained by both an increase of within-category correlations (average slope of linear fit across layers = 0.046; p = 0; Supplementary Figure 1, top left), and a decrease of between-category correlations across layers (average slope across layers = −0.008; p = 0; Supplementary Figure 1, top right).

**Figure 3:**
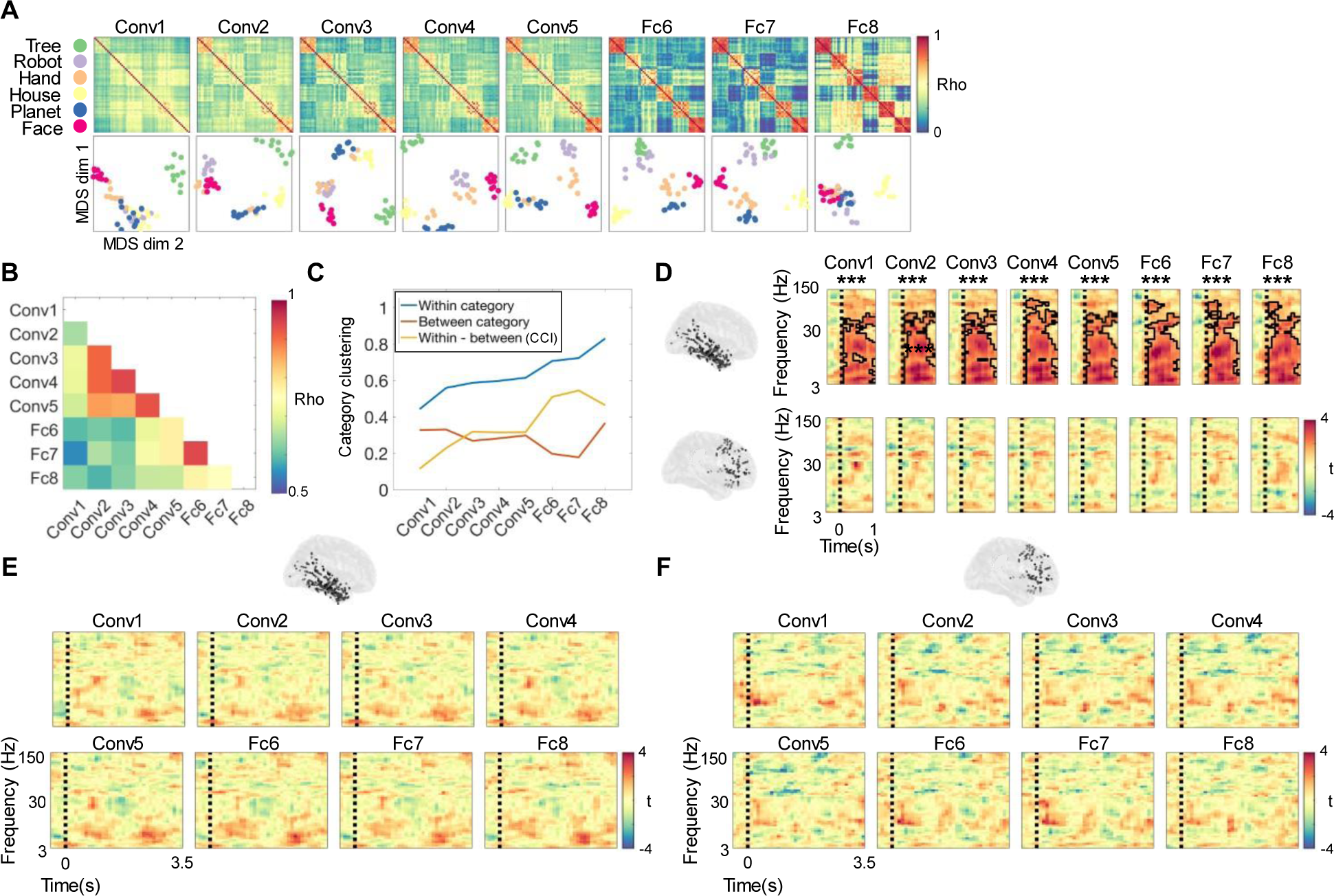
Analysis of representational formats using a feedforward deep neural network. (A) Top: Representations in the feedforward network AlexNet. Representational Similarity Matrices (RSMs) reflecting pairwise correlations of unit activations in each layer of the network. Bottom: 2D multidimensional scaling (MDS) projections of RSMs at each layer, color-coded according to categories. (B) Representational consistency plot showing pairwise correlations (Spearman’s rho) of RSMs at each network layer. (C) Within-category, between-category and within-category vs. between-category correlations (i.e., Category Cluster Index, CCI) as a function of network layer. (D) Top: Correlations between RSMs from the DNN and neural data, for each AlexNet layer and each encoding time-frequency window in the VVS. Each time-frequency plot shows the correlation values of representations in one particular layer to neural representations. Clusters outlined in black indicate time-frequency periods where correlation values are significantly higher than zero at the group level (Bonferroni corrected for 8 layers). Bottom: Same analysis for PFC data. (E) No matching of VVS RSMs with AlexNet RSMs during the maintenance period. (F) Same analysis as in E for the PFC data.

We next set out to evaluate the similarity between stimulus representations in AlexNet and neural representations in VVS and PFC. In order to characterize the frequency profile of reactivations, we performed a frequency-resolved analysis of fits between neural and DNN representations: We constructed RSM time-series for every frequency independently and grouped them into a time-frequency map of model fits (Methods). In the VVS, we found that representational geometries during encoding were captured by network representations in all layers in the 3-75Hz range (all p_corr_ < 0.008); Figure 3D, top row). In layers 4 and 6-8, this effect extended into the high gamma frequency range. Similar to the results observed in the category model analysis (Figure 2B), we did not observe any matching between neural and AlexNet representations during the maintenance period (all p_corr_ > 0.056; Figure 3E). In the PFC, we did not observe any significant fit during either encoding (all p > 0.064; Figure 3D, bottom row) or maintenance (all p > 0.168; Figure 3F).

Taken together, these results show that representations in the AlexNet are aligned with encoding representations in VVS but not PFC. Importantly, during the maintenance period neither VVS or PFC representations showed a significant fit with representations in the AlexNet network, suggesting that the format of prioritized VWM representations cannot be explained by feedforward DNNs.

We thus employed the recurrent neural networks BL-NET and corNET-RT to characterize representational formats in VVS and PFC. We first assessed the temporal evolution of network representations in the different layers of BL-NET and correlated the layer-wise RSMs between successive time points (Methods). In all layers, representations changed most prominently between intervals 1-2 and least between intervals 7-8 (Figure 4A). In layers 2 to 7, representations remained largely constant following time step 3, while the first layer showed more substantial dynamics until the last time interval (Figure 4A). Directly comparing the representations between the initial (1^st^) and the final (8^th^) time points separately for each layer revealed larger changes in the first two layers and substantially smaller changes in layers 3-7 (Figure 4B and 4C). Similar to the AlexNet, CCI values were significantly higher than chance in all layers (all p = 0.007, Bonferroni corrected for 7 layers), and we observed a fourfold increase of CCI values from the first (CCI = 0.07) to the last layer (CCI = 0.40) of the network (Figure 4D). Contrary to the AlexNet, the increases in CCI in the BL-NET network were only due to a decrease in between-category correlations (average slope of linear fit across layers = −0.06; p = 0; Supplementary Figure 1, middle right), while the within-category correlations did not change across layers (average slope of linear fit across layers = −0.00062; p = 0.72; Supplementary Figure 1, middle left). Furthermore, BL-NET between-category correlations decreased significantly more across layers than AlexNet between-category correlations (Alexnet vs. BL-NET slope difference = 0.1061; p = 0).

**Figure 4:**
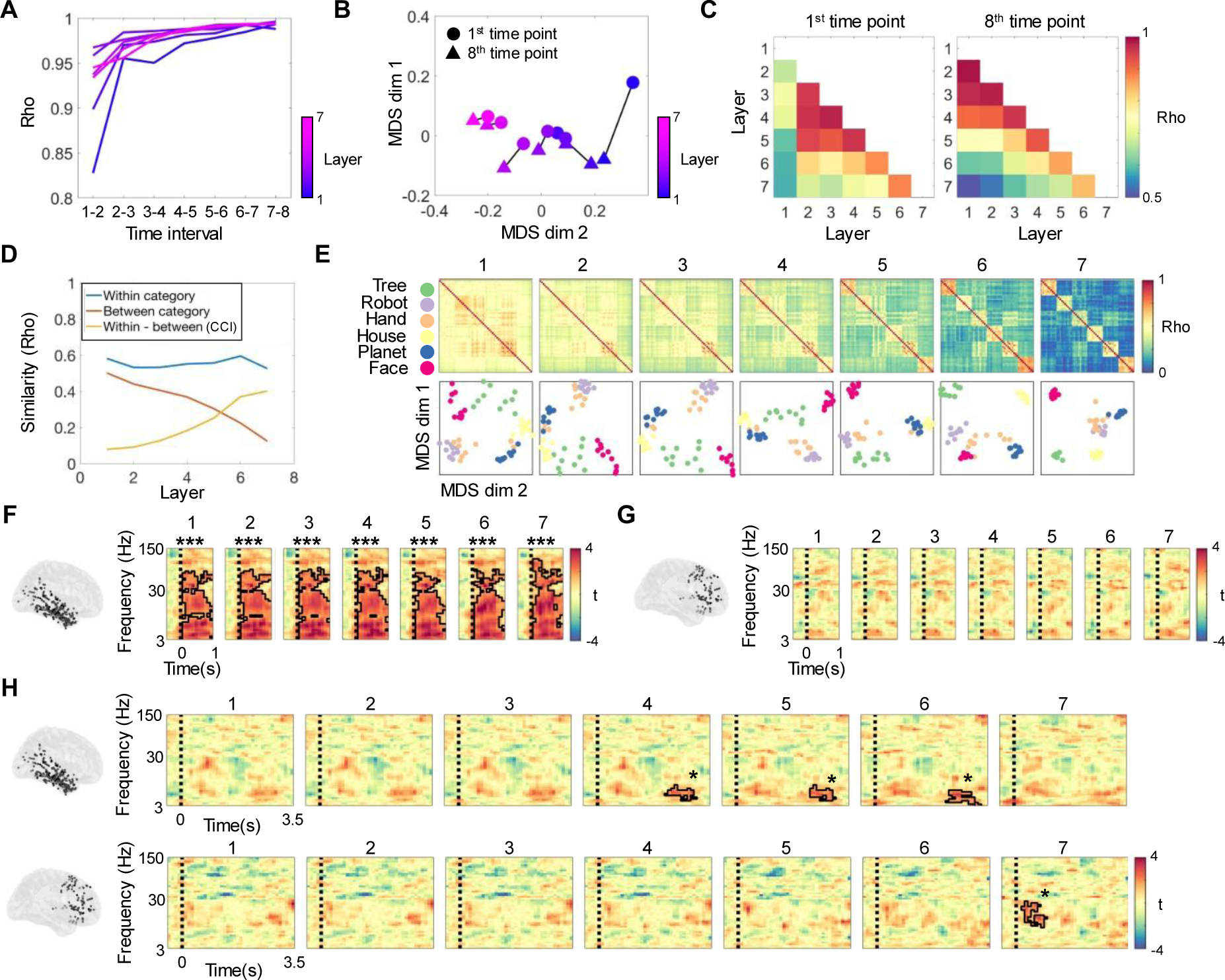
Analysis of representational formats using the BL-NET network. (A) Representational consistency at each time interval of the BL-NET network was computed by correlating representations formed at successive time points. Each curve represents one layer of the network, color-coded from early (blue) to deep layers (pink). (B) Two-dimensional projections of the first and last time point of each layer in the BL-NET network showing greater representational distances in the first layer than in all other layers. (C) Pairwise correlations of RSMs corresponding to the first (left) and last (right) time points in each layer of BL-NET. (D) Within-category, between-category and within-category vs. between-category correlations (i.e., Category Cluster Index, CCI) for each layer of BL-NET (last time point). (E) RSMs and corresponding MDS projections for the last time point of all BL-NET layers. In the MDS plots, items are color-coded according to category. (F) Correlations between BL-NET RSMs (last time point in each layer) and neural RSMs during encoding in VVS. Outlined clusters indicate time-frequency periods where correlation values are significantly higher than zero at the group level (Bonferroni corrected for 7 layers). (G) Same analysis as in F for the PFC. (H) Same analysis as in F for the maintenance period in the VVS (top) and PFC (bottom). In the VVS, BL-NET representations in layers 4, 5, and 6 matched representations in the theta/alpha frequency range (3-14Hz) prior to the probe. In the PFC, BL-NET representations in the last layer matched representations in the beta band (16-29Hz) following presentation of the retro-cue. ***: p < 0.001.

Together, these results show that the BL-NET network represents low-level features more dynamically than abstract high-level features and that it clusters categorical information more strongly in deep than superficial layers. Contrary to the AlexNet network, this clustering is exclusively due to a reduction of between-category correlations rather than an increase in within category correlations across network layers.

We next compared neural and BL-NET representations, focusing on the RSMs at the last time-point of each layer (Figure 4E). During encoding, results were similar to those in the AlexNet analysis: Network representations of all layers matched VVS representations for a wide range of frequencies between 3 and 75Hz (p_corr_ = 0.007; Figure 4F), and these extended into the high-gamma range (i.e., until 110Hz) in layer 7. No significant correlations were observed in the PFC (all p_corr_ > 0.263; Figure 4G). During maintenance, no significant fits were observed in the VVS following the retro-cue, again consistent with the AlexNet analysis. Interestingly, however, we observed a significant matching of VVS representations in the theta/alpha frequency range (3-14Hz) with BL-NET representations in layers 4 (p_corr_ = 0.035), 5 (p_corr_ = 0.035) and 6 (p_corr_ =0.014). These effects occurred in a late maintenance time period from 2.1s to 3.2s, close to the presentation of the probe (Figure 4H, top row). Critically, in the PFC, we observed a significant fit between neural and network RSMs following presentation of the retro-cue, i.e. time-locked to the prioritization process. This effect started 200ms after the onset of the retro-cue and lasted for 800ms; It was specifically observed for the last layer of the BL-NET (final layer: p_corr_ = 0.021; all other layers: p_corr_ > 0.43), and related to neural representations in the beta frequency range (15-29Hz; Figure 4H, bottom row).

To verify that these results were not driven by the particular dataset with which the BL-NET was trained (i.e., ImageNet, as in previous studies: (Baek et al., 2021; Bao et al., 2020; Cadieu et al., 2014; Cichy et al., 2016; Khaligh-Razavi & Kriegeskorte, 2014; Kuzovkin et al., 2018; Liu et al., 2020, 2021; H. Tang et al., 2018; Vinken & Op de Beeck, 2021), we performed the same analysis with a variant of this network trained with a recently released dataset of images (Mehrer et al., 2021). The ‘Ecoset’ dataset includes >1.5 million images from 565 categories which were selected to capture the distribution of objects relevant to humans in natural conditions, thus providing an ecologically valid alternative to ImageNet (Methods). We computed BL-NET fits for both the VVS and the PFC during encoding and maintenance employing the Ecoset-trained network weights. We focused on the specific time-frequency clusters where we observed significant effects in the BL-NET analysis using the ImageNet training set.

Ecoset-trained BL-NET representations during encoding significantly matched VVS representations in the encoding clusters observed before (all p < 0.0001). During maintenance, results were also significant for clusters observed in layers 4 (p = 0.0003), 5 (p = 0.0005) and 6 (p = 0.0001). In the PFC, we observed a significant correlation with representations in the last layer of the Ecoset-trained BL-NET (p = 0.002), again confirming the abstract representational format of PFC representations following the retro-cue.

In our final analysis, we employed the corNET-RT model to account for VVS and PFC representations. Consistent with the BL-NET analyses, we first evaluated the representational consistency across successive time points in each layer of the network. The final layer (IT) showed the lowest correlation across consecutive time points compared to all other recurrent passes in the network (Rho = 0.78; note the first recurrent pass in IT is the fourth overall pass in the network, Methods). This demonstrates that contrary to the BL-NET, corNET-RT represents stimuli more dynamically in its deepest layer. In addition, we observed that representations in layers V2 and V4 clustered together in representational space, while representations in V1 and IT were segregated (Figure 5B and 5C). Categorical clustering of representations was found in all layers, as evidenced by significant CCI scores in each layer and at each time point (all p_corr_ > 0.004; Figure 5D). Similar to BL-NET and contrary to AlexNet, we observed a prominent increase in CCI scores across layers, which was mostly due to a decrease in between-category correlations (average slope across layers = −0.14; p = 0; Figure 5D and Supplementary Figure 1). However, within category correlations were also reduced across network layers (average slope across layers = -0.02; p = 1.6301e-11; Figure 5D and Supplementary Figure 1).

**Figure 5:**
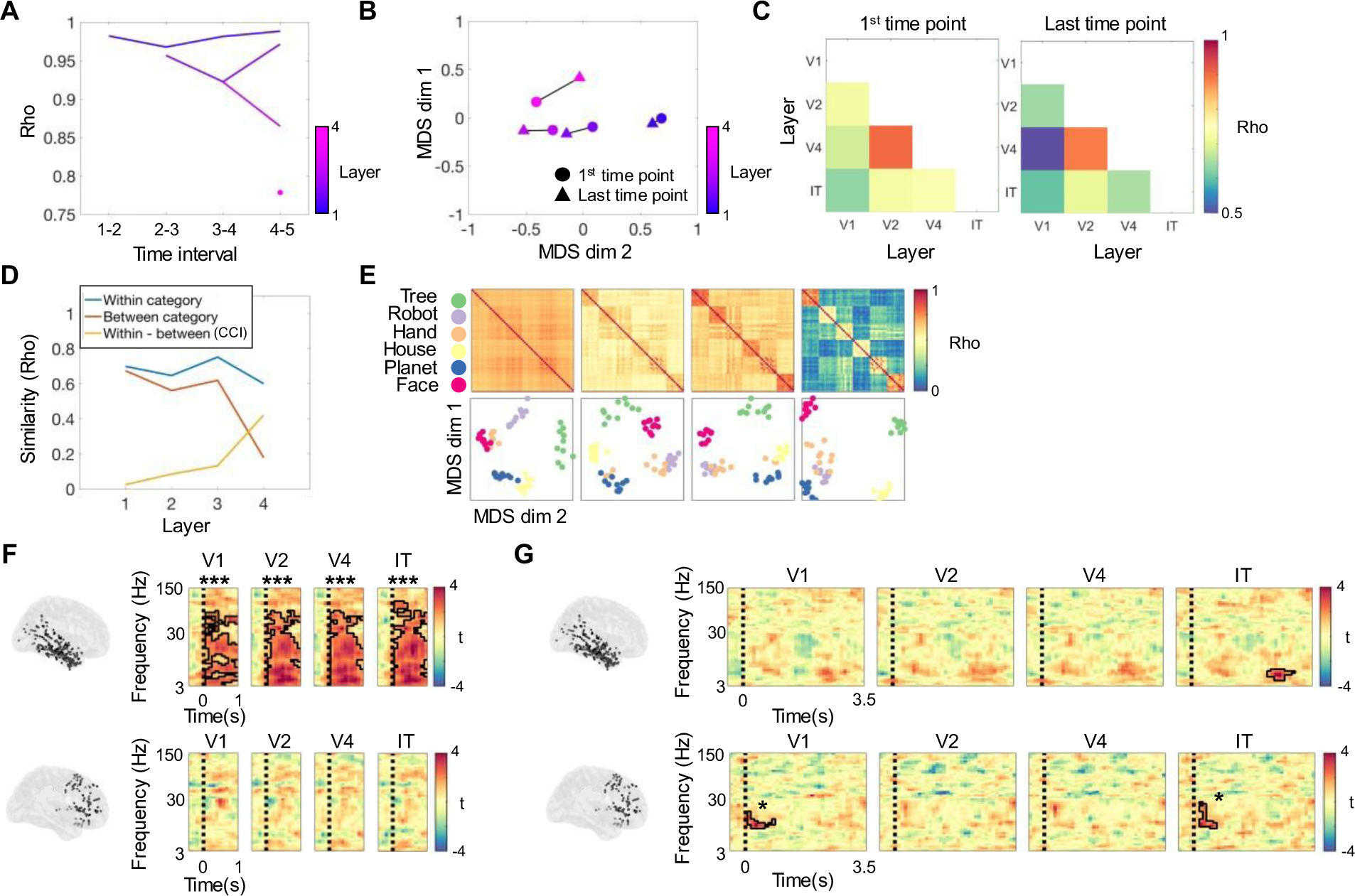
Analysis of representational formats using the corNET-RT network. (A) Representational consistency at each time interval of the corNET-RT network. Note that in this architecture, different layers have different numbers of recurrent passes, and deep layers do not receive input until activity has propagated from early layers. Each curve represents one layer, color-coded from early (blue) to deep (pink). (B) MDS projections of first (circle) and last (triangle) time point in each layer show relatively higher temporal dynamics in layer IT (output) compared to the other layers. MDS results have been scaled for visualization. (C) Pairwise correlations of RSMs corresponding to the first (left) and last (right) time points in each layer of corNET-RT. (D) Within-category, between-category and within-category vs. between-category correlations (i.e., Category Cluster Index, CCI) at each layer of the network (final time point). (E) RSMs (top) and corresponding MDS projections (bottom) for each layer of the corNET-RT network (final time-point). Items are color-coded by category. (F) Correlations between corNET-RT RSMs (last time point in each layer) and neural RSMs during encoding in VVS (top) in the 3-150Hz frequency range. Colors indicate resulting t-maps in the comparison of group-level correlation values against zero. Significant regions after multiple comparisons correction are outlined in black (Bonferroni corrected for 4 layers). (G) Same analysis as in F for the maintenance period. Top: A match between network and neural representations was observed in the VVS in a late period, close to the presentation of the probe, in the IT layer. Bottom: In the PFC, correlations were significant with representations in the beta frequency range (15-29Hz) following the onset of the retro-cue with both V1 and IT layer. *** = p < 0.001; * = p < 0.05

We compared corNET-RT RSMs to neural representations, focusing on the last time point in each layer, again consistent with the BL-NET analysis (Figure 5E). During encoding, we found a significant match of VVS representations across a wide range of frequencies with corNET-RT representations in all layers (3-105Hz; all p_corr_ < 0.004; Figure 5F, top row). No significant correlations were found in the PFC (all p_corr_ > 0.053, Figure 5F, bottom row). During the maintenance phase, corNET-RT representations in IT matched those in the VVS towards the end of the maintenance period, specifically in the theta-alpha frequency range, consistent with the results observed in the BL-NET analysis (6-11Hz; p_corr_ = 0.044; Figure 5G, top row). Critically, we again observed a significant match of corNET-RT representations in IT with PFC representations time-locked to the presentation of the retro-cue and in the beta band (15-29Hz; p_corr_ = 0.016; Figure 5I, right). This effect lasted for 500ms, similar to the results observed in the BL-NET analysis. In addition, we observed a significant correlation with representations in V1 (p_corr_ = 0.036).

Taken together, these results show that PFC representations following the retro-cue matched those in two recurrent neural network architectures (the BL-NET and the corNET-RT) but not those of a purely feedforward network (the Alexnet), and that these effects were specific to the beta-frequency range and most prominent for late layers of the networks. VVS representations did not show correspondence with representations in recurrent networks following the retro-cue, but prior to the probe.

## Discussion

Our study aimed to unravel representational formats and neural coding schemes in sensory and executive control regions during WM prioritization. Specifically, we analyzed the impact of WM prioritization on stimulus-specific activity patterns in VVS and PFC and assessed their representational formats using feedforward and recurrent DNN models of natural image processing. The VVS exhibited pronounced category-specific representations during encoding which were reinstated during the maintenance period, reflecting a shared (or ‘mnemonic’) coding scheme across both experimental phases. The PFC exhibited robust category-specific representations during WM encoding as well, but did not show reinstatement of encoding patterns during the maintenance period. Subsequent in-depth analyses showed that this lack of reinstatement in PFC was not due to memory decay or reduced signal to noise ratio, but due to a transformation of representations between different task-dependent formats, in line with a dynamic ‘prioritization’ coding scheme: Representations in PFC corresponded to a simple categorical model during encoding, but matched only high-level visual and abstract formats from a recurrent DNN following the retro-cue. This shift was also reflected at the level of the neurophysiological substrates of WM representations, since PFC representations during encoding were observed in theta, beta and gamma frequency bands but exclusively in beta frequency oscillations during the retro-cue. Taken together, these results demonstrate that WM prioritization relies on a distinct recruitment of specific task-depend representational formats in the PFC.

Recent investigations showed a transformation of visual representations from perceptual to abstract formats during VWM encoding (Liu et al., 2020, 2021). While representations in these studies were based on patterns across the entire brain, we here focused on representations in two brain regions that are critical for VWM storage and control, respectively: VVS and PFC. We note that our initial RSA analysis of category representations (Figure 2A) could not explain representations during the maintenance period in either of these regions. The EMS analysis (Figure 2F), however, revealed a distinct set of results in VVS and PFC: While encoding activity patterns reoccurred during the maintenance period in the VVS, this was not the case in the PFC. In the VVS, representations towards the end of the maintenance period matched representations in intermediate and deep layers of two recurrent DNN architectures (BL-NET and corNET-RT), suggesting a transformation during encoding from a purely ‘categorical’ format into a format that incorporates high-level visual and semantic relationships among stimuli. Thus, despite a relative stability of neural activity patterns (as revealed by the EMS analysis), their representational geometry changes and eventually results in less categorical representations during the maintenance period.

By contrast, in the PFC, encoding activity patterns did not reoccur during the maintenance period, suggesting a more pronounced transformation in this region; however, neural representations following the retro-cue matched representations in deep layers of two recurrent DNN architectures. Our results thus provide evidence for two distinct representational transformations which are both critically reliant on deep recurrent computations: (1) In VVS, from a purely categorical format into a format that is subsequently reinstated during maintenance; (2) in PFC, from a categorical format during encoding into a different format that occurs only following the retro-cue. Since representations in the VVS are shared between encoding and maintenance, we refer to them as a ‘mnemonic’ coding scheme; in contrast, the distinct representational format in PFC following the retro-cue corresponds to a ‘prioritization’ coding scheme.

What could be the functional role of the transformation of category-specific representations in the PFC? Our data is consistent with a capacity-limited view of WM which would benefit from compressed stimulus representations in this region. This notion has been recently supported by behavioral (Kerren et al., 2022) and neuroimaging (Kwak & Curtis, 2022) studies. In particular, (Kerren et al., 2022) demonstrated that semantic aspects of images are selectively prioritized during WM maintenance in a multi-item WM task, while no selective storage of abstract features of images is present in single-item tests. This fits to our findings in the VVS that contained both perceptually detailed and abstract representations during maintenance of single items, while we additionally found abstract representations in the PFC. In the fMRI study of (Kwak & Curtis, 2022), representational abstraction was observed in parietal and visual cortices, but not in prefrontal regions. The differences between our results and those of (Kwak & Curtis, 2022) might relate to the particular stimuli employed (natural images with semantic content versus low-level visual features), and to our use of a paradigm involving prioritization, which preferentially engages the PFC (Myers et al., 2017; Wallis et al., 2015).

Abstract information in PFC was specifically detected in the 15-29Hz frequency range, i.e., within the beta band (13-29Hz). Previous studies showed a prominent role of prefrontal beta oscillations for top-down control of information in WM (Lundqvist et al., 2018; Miller et al., 2018), and oscillatory activity in the beta range has also been associated with transient task-dependent activation of stimulus-specific information during WM maintenance (Spitzer & Haegens, 2017). Our results in PFC are well consistent with these interpretations: The representations we observed are content-specific and locked to the presentation of the retro-cue, which is when the prioritization process takes place. This aligns with previous studies that have reported stimulus-specific activity in the beta range during WM maintenance (e.g., (Wimmer et al., 2016); see (Spitzer & Haegens, 2017) for review). In contrast to standard delay tasks where beta modulations occur late in the WM delay period (Spitzer et al., 2010; Wimmer et al., 2016), our study demonstrates brief and cue-locked effects, consistent with previous retro-cue paradigms (Spitzer & Blankenburg, 2011). Our findings are also consistent with the widely accepted role of the PFC in the top-down control of information stored in other brain regions, in line with previous studies on both episodic and working memory (Eichenbaum, 2017). Indeed, activity in the PFC has been linked to task-dependent executive control over specific contents in several studies (for a review, see Rissman & Wagner, 2012). This could be achieved by modulating the activation state of distributed perceptual and mnemonic representations (Eichenbaum, 2017), for instance through PFC connectivity with the VVS (Ten Oever et al., 2021). The beta-frequency reactivation we observed in the PFC is suggestive of a top-down signal prompted by presentation of the cue that might affect information processing in downstream regions. Further studies are required to investigate this possibility. Nevertheless, our results confirm previously untested views of PFC functioning by demonstrating its engagement in abstract VWM representations during VWM prioritization.

DNNs are increasingly used in cognitive neuroscience to characterize the representational formats and temporal dynamics of perceptual and mnemonic representations in the human brain. While different feedforward and recurrent architectures have been applied in the domain of vision, resulting in a wide variety of models employed to fit neural data (e.g., Cadieu et al., 2014; Kar et al., 2019; Kar & DiCarlo, 2021; Khaligh-Razavi & Kriegeskorte, 2014; Kietzmann et al., 2019; Yamins et al., 2014), this approach has only started to be employed in memory research. Pioneering investigations have applied the feedforward neural network AlexNet to study representational formats during visual working memory in humans (Liu et al., 2020, 2021). Notably, these studies did not investigate the representational formats during WM maintenance but focused solely on the encoding period. While theoretical and experimental considerations have strongly argued for the use of recurrent architectures in the domain of visual perception (Kietzmann et al., 2019; van Bergen & Kriegeskorte, 2020; Yamins & DiCarlo, 2016), they have so far not been applied to memory research. The use of recurrent architectures in the context of working memory is particularly important given the relevance of recurrent computations for PFC processing (Mante et al., 2013; Miller & Cohen, 2001) and WM functions in general (Bouchacourt & Buschman, 2019; Ehrlich & Murray, 2022; Gelastopoulos et al., 2019; Wan et al., 2022). In our study, we tested a feedforward and two recurrent models in their ability to predict representational distances in human iEEG data. During encoding, both types of models captured the representational geometry of stimuli across all layers and a wide range of frequencies in the VVS, while no fits were observed in the PFC (Figure 3, 4 and 5). During maintenance, however, the two architectural families strongly differed in their fit to the neural data: the AlexNet was unable to capture representations in either region, while BL-NET and corNET-RT matched representations in both VVS and PFC (Figure 4I and 5I). Together, these results demonstrate that only recurrent architectures can explain the representational geometry of stimuli during VWM prioritization, while a feedforward architecture and a simple model of category information do not provide good fits.

What are the differences in the representational geometries of Alexnet, BL-NET and corNET-RT that can explain the different fits to the neural data we observed? We thoroughly characterized within-category, between-category and within-vs. between-category correlations (i.e., CCI) in all three architectures to investigate their differences in stimuli representation. We found that all networks represented increasingly category-specific information across layers, as assessed by prominent increases in CCI, yet this was achieved through different representational changes. While the AlexNet showed both an increase in within-category correlations and a decrease in between-category correlations, the recurrent models only showed decreases in between-category correlations (Supplementary Figure 2), suggesting that recurrence particularly supports distinct representations of different categories. Again, further studies are needed to unravel the possible neurophysiological basis and cognitive function of these representational transformations.

Computational models of WM have proposed that prioritization requires a transformation of the neural space of activity in which the items are represented, involving, for example, a rotation or “flip” of the format of prioritized content in neural activity space (Wan et al., 2022). These models have recently received empirical support from studies in monkeys (Panichello & Buschman, 2021) suggesting an efficient neural code that organizes and structures neural representations during the prioritization process. Consistently, a recent iEEG study in humans demonstrated a role of PFC in resolving cognitive interference between competing sensory features by transforming their representational population geometry into distinct neural subspaces to accommodate flexible task-switching (Weber et al., 2023). Our work contributes to this literature by establishing that the PFC not only supports a transformation of the representational geometry of stimuli but also a differential representation of particular visual formats in the context of VWM prioritization.

It has been recently proposed that the selection of network training sets critically influence the matching of DNN and neural representations, and that this influence may be more important than specific architectural constraints (Conwell et al., 2023). For this reason, in our study we employed two different datasets of images (Imagenet and Ecoset), which provided largely consistent results (Supplementary Figure 2). Other limitations remain, however: First, the BL-NET and corNET-RT networks were not trained to memorize stimuli but to solve the task of image classification, which may be argued to limit their heuristic value as models of WM representation. It is interesting to see that despite this different objective function they significantly match the representations during WM prioritization. Second, these models do not include top-down connections but only lateral connectivity, and thus cannot account for PFC-VVS interactions. In future work, novel architectures should be employed that mimic brain connectivity more accurately at least at a high-level of description (i.e., containing top-down as well as within-layer connections). Finally, while we decided to focus on the prioritization process and the single-item trials in this study, we aim to further investigate the representation of multiple items in the future. A promising avenue for this purpose is the use of sequential recurrent convolutional networks that receive multiple consecutive images as input and can be employed to track multi-item representations (Sörensen et al., 2023).

In summary, we present evidence of successive representational transformations during VWM encoding and after item prioritization in the VVS and the PFC that critically depend on recurrent computations and abstract representational formats. This result shows that percepts originally formed during encoding are differentially abstracted and reshaped in VVS and PFC to enable flexible task-dependent manipulations during working memory prioritization.

**Supplementary Figure 1:**
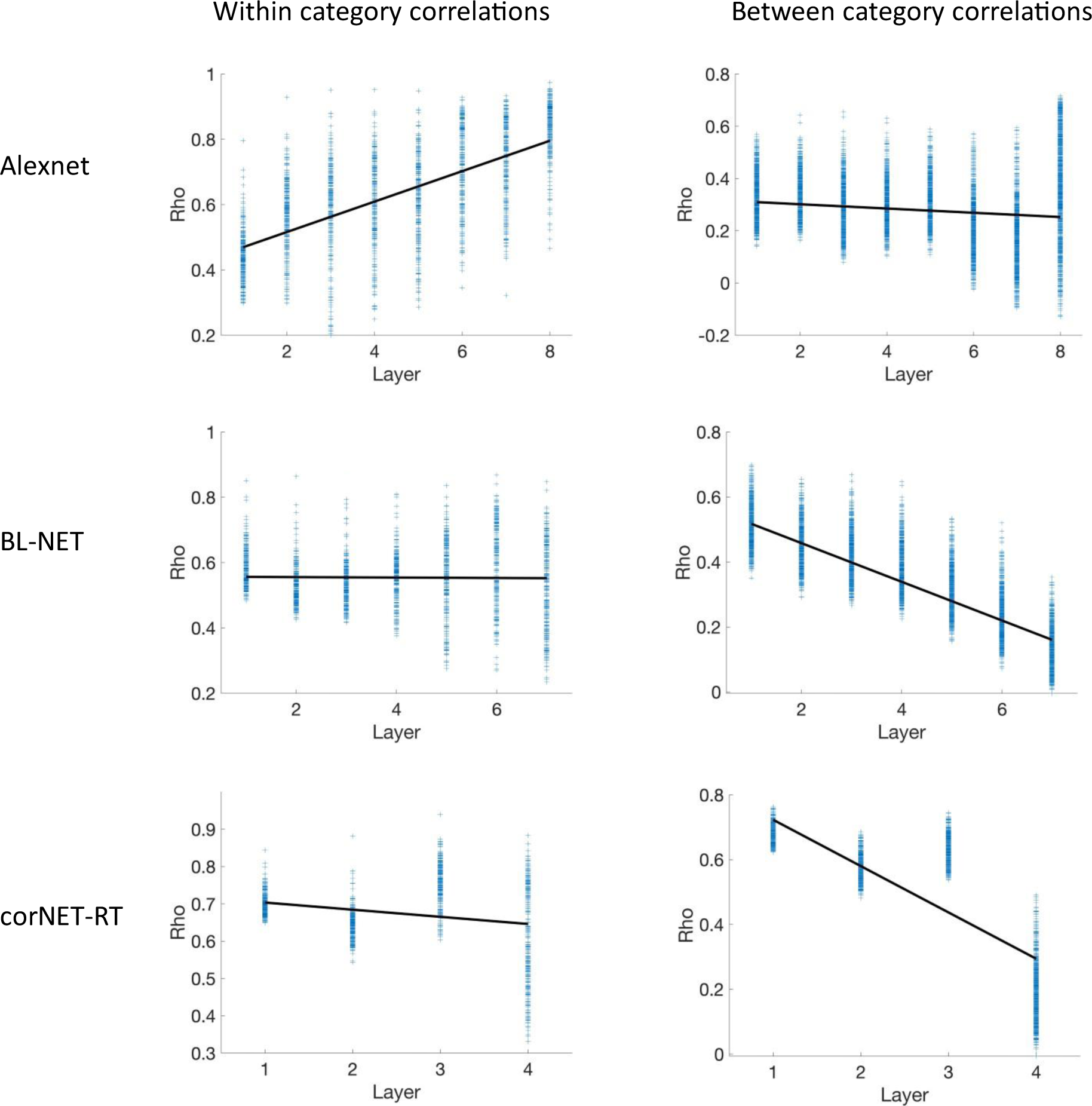
Within-category and between-category correlations as a function of network layer for all employed DNNs. The figure shows all pairwise within-category (left column) and between-category (right column) correlations for AlexNet (top row), BL-NET (middle row), and corNET-RT (bottom row) networks. Black line in each plot shows linear fit performed over all pairwise comparisons in each network for visualization, but note that the statistics reported in the main text were computed by performing a linear fit to each of the individual item pairs separately, and contrasting the resulting distribution of slopes across networks and conditions.

## Methods

### Participants

Thirty-two patients (17 females, 30 ± 10.04 years) with medically intractable epilepsy participated in the study. Data were collected at the Freiburg Epilepsy Center, Freiburg, Germany; the Epilepsy center, Second Affiliated Hospital, School of Medicine, Zhejiang University, Hangzhou, China; and the Center of Epileptology, Xuanwu Hospital, Capital Medical University, Beijing, China. The study was conducted according to the latest version of the Declaration of Helsinki and approved by the responsible ethics committees, and all patients provided written informed consent.

### Experimental design

Participants performed a multi-item working memory paradigm involving a retro-cue. They encoded a sequence of 3 images of natural objects from different categories and were asked to remember this content during a subsequent maintenance period. This period consisted of two phases that were separated by a retro-cue. The retro-cue prompted participants to selectively maintain items from particular encoding positions (single-item trials, 50%), or to maintain all items in their order of presentation (multi-item trials, 50%) for a subsequent memory test. Note that with the exception of the behavioral data, we only focused on the single-item trials in this study. In the test, six items were presented, which included all 3 presented items from encoding, two new exemplars from two of the encoding categories, and 1 item from a different category (Figure 1A). In the single-item trials, one of the lure items in the test was always from the same category as the cued item. Objects pertained to six categories (trees, robots, hands, houses, planets and faces) with ten exemplars each (60 images in total; Figure 1B). In order to perform the task correctly, participants needed to remember not only categorical information about the items but also the specific perceptual information identifying each individual exemplar.

In the multi-item trials, we considered as correct sequences in which all items were correctly retrieved in the right order. Note that performance for single and multi-item trials have by definition different chance levels. We computed chance levels through simulations with the real randomized sequences given to our participants. We designed an “agent” that performed the task randomly (i.e., selecting an option randomly from the available options in each trial). We run the simulations 1000 times and averaged performance for item and category levels. We confirmed that in the single-item trials, chance levels in our simulations approximated the real chance values: 0.167 ± 0.001 (performance expected by chance: 0.167). In the multi-item trials, chance levels were at 0.0083 ± 0.00052.

The task was divided into blocks and sessions. Each block consisted of 60 trials. Each session consisted of at least one block, but most participants performed between 1 and 3 blocks in each session (2.59 ± 1.04) and between 1 and 2 sessions (1.19 ± 0.39) in total. The order and frequency of image presentations was pseudorandomized to balance repetitions of images across blocks and sessions. The experiment was programmed in Presentation (Neurobehavioral systems, California, USA), and was deployed on Samsung 12’’ tablet computers running Microsoft Windows. Patients performed the experiment while sitting in their hospital beds and responded to the memory test utilizing the touch-screen of the tablet.

Two versions of the experiment were implemented for the different patient populations in Germany and China. The two versions had identical stimuli in all categories except for the category “ aces”. The German version of the experiment included faces of former German chancellor Angela Merkel and the Chinese version included faces of the actor Jackie Chan. This was made to ensure that the face represented was equally well known to the different patient populations.

### Intracranial EEG recordings

IEEG data were recorded using amplifiers from Compumedics (Compumedics, Abbotsford, Victoria, Australia), and Brain Products GmbH with sampling rates of 2,000Hz and 2,500Hz, respectively. Patients were surgically implanted with intracranial depth electrodes for seizure monitoring and potential subsequent surgical resection. The exact electrode numbers and implantation locations varied across patients and were determined by clinical needs. Online recording data was referenced to a common scalp reference contact which was simultaneously recorded with the depth electrodes. Data was downsampled to 1,000Hz and bipolarized by subtracting the activity of one contact point with that from the nearest contact of the same electrode, resulting in a total of N-1 virtual channels for an electrode with N channels after bipolarization.

### Channel Localization

Electrodes employed were standard depth electrodes (Ad-Tech Medical Instrument Corporation, Winsconsin, USA). Electrodes contained variable number of contacts and inter-contact distances. In the data collected at Zhejiang University, Hangzhou, and Medical University, Beijing, each depth electrode was 0.8 mm in diameter and had either 8, 12 or 16 contacts (channels) that were 1.5cm apart, with a contact length of 2 mm. Channel locations were identified by coregistering the post-implantation computed tomography (CT) images to the pre-implantation Magnetic Resonance Images (MRIs) acquired for each patient, which were afterwards normalized to MNI space using Statistical Parametric Mapping (SPM; https://www.fil.ion.ucl.ac.uk/spm/). We then determined the location of all electrode channels in MNI space using PyLocator (http://pylocator.thorstenkranz.de/), 3DSlicer (https://www.slicer.org) and FreeSurfer (http://surfer.nmr.mgh.harvard.edu). In a group of patients (data collected in Beijing), we determined MNI coordinates using the pipeline described in (Stolk et al., 2018), and identified the closest cortical or subcortical label for each channel in each patient. In all patients, we removed channels located in white matter, resulting in 588 clean channels across all patients (18.4 ± 11.9 channels per patient).

### ROI selection

We selected two main regions of interest given their well-known involvement in VWM: the ventral visual stream (VVS) and the prefrontal cortex (PFC). The VVS has been widely studied in the context of object recognition during visual perception (Cichy et al., 2014). Previous work employing iEEG and Deep Neural Networks often applied RSA metrics to activity from distributed electrodes across the whole brain (e.g., Liu et al., 2020), and we specifically aimed to extend these studies by investigating region-specific representations in the context of VWM (see also (Ten Oever et al., 2021).

The role of the PFC in working memory has been linked to executive control processes that enable the task-dependent manipulation and transformation of information (D’Esposito et al., 1995; Miller et al., 2018; Myers et al., 2017). However, relatively little is known about the representational formats of VWM representations in this region during prioritization. Moreover, no previous study investigated region-specific representational similarity during VWM.

Electrodes located at the following freesurfer locations were labeled as S electrodes ‘inferior temporal’, ‘middle temporal’, ‘superior temporal’, ‘bankssts’, ‘fusiform’, ‘cuneus’, ‘entorhinal’. Electrodes with the following labels were categorized as PFC electrodes ‘medial orbitofrontal’, ‘pars triangularis’, ‘superior frontal’, ‘lateral orbitofrontal’, ‘pars opercularis’, ‘rostral anterior cingulate’, ‘rostral middle frontal’, ‘superior frontal’. Electrodes from both left and right hemispheres were included in our ROIs.

### Preprocessing

We visually inspected raw traces from all channels in each subject independently and removed noisy segments without any knowledge about the experimental events / conditions. All channels located within the epileptic seizure onset zone or severely contaminated by epileptiform activity were removed from further analyses. We divided the data into 9-second epochs (from -2 to 7s) around the presentation of each stimulus at encoding or the onset of the retro-cue during the maintenance period. After epoching the data, we completely removed epochs containing artifacts that were identified and marked in the non-epoched (continuous) data. We visually plotted spectrograms to verify the presence of artifacts in the frequency domain in the resulting epochs. The number of epochs corresponding to item or cue presentation that were removed varied depending on the quality of the signal in each subject (15.10 ± 13.14 in each session, corresponding to around 6.9% of all epochs in each session).

Preprocessing was performed on the entire raw data using EEGLAB (Delorme & Makeig, 2004), and included high-pass filtering at a frequency of 0.1Hz and low-pass filtering at a frequency of 200Hz. We also applied a band-stop (notch) filter with frequencies of 49-51Hz, 99-101Hz, and 149-151Hz.

### Time-frequency analysis

Using the FieldTrip toolbox (Oostenveld et al., 2011), we decomposed the signal using complex Morlet wavelets with a variable number of cycles, i.e., linearly increasing in 29 steps between 3 cycles (at 3 Hz) and 6 cycles (at 2 Hz) for the low-frequency range, and in 25 steps from 6 cycles (at 30 Hz) to 12 cycles (at 150 Hz) for the high-frequency range. These time-frequency decomposition parameters were taken following previous research that used iEEG oscillatory power as features for RSA (Liu et al., 2020; Pacheco Estefan et al., 2019; Staresina et al., 2016). The resulting time-series of frequency-specific power values were then z-scored by taking as a reference the mean activity across all trials within an individual session (Fellner et al., 2020). This type of normalization was applied to remove any common feature of the signal unrelated to the encoding of stimulus-specific information. We z-scored across trials in individual sessions in our final analyses, but similar results were obtained when we z-scored the data by considering the activity of all trials irrespective of the session. We employed the resulting time-frequency data to build representational feature vectors in our pattern similarity analyses (see below).

### Pattern similarity analysis: representational patterns

We employed different representational features in our analyses involving model RSMs (i.e., the category model RSA analysis and the DNN-based RSA analyses), and our analyses involving particular contrasts (i.e., encoding-encoding similarity analysis [EES] and the encoding-maintenance similarity analysis [EMS]; see below). In both types of analyses, representational feature vectors were defined by specifying a 500ms time window in which we included the time courses of frequency-specific power values in time-steps of 100ms (5 time points) across all contacts in the respective ROI (VVS or PFC). In the RSM based analyses, we performed this analysis separately for each individual frequency in the 3-150Hz range, while in the EES and EMS analyses, we analyzed activity patterns across individual frequencies within five different bands (theta, 3-8Hz; alpha, 9-12Hz; beta, 13-29Hz; low-gamma, 30-75Hz, high-gamma, 75-150Hz). In the RSM-based analyses, a frequency-specific representational pattern was thus composed of activity of N electrodes x 5 time-points in each 500ms window. In the EES and EMS analyses, this representational feature vector consisted of N electrodes x M frequencies x 5 time-points. Note that the number of channels included in the representational feature vectors varied depending on the number of electrodes available for a particular subject/ROI, and the number of frequencies included in each band in the EES and EMS analyses varied as well (theta: 6 frequencies; alpha: 4 frequencies; beta: 17 frequencies; low-gamma: 9 frequencies, high-gamma: 16 frequencies; see section *Time-frequency decomposition* above). These two- or three-dimensional arrays were concatenated into 1D vectors for similarity comparisons. Only subjects with at least 2 electrodes in a particular ROI were included in all RSA analyses, leading to 15 subjects in the PFC and 26 subjects in the VVS.

### Model-based RSA

We employed temporally resolved Representational Similarity Analysis (RSA) to evaluate the dynamics of categorical information in our data following previous work (Cichy et al., 2014; Kietzmann et al., 2019). A main assumption of this research is that stimuli from the same categories will have greater neural similarity than stimuli from different categories. To evaluate this hypothesis, we constructed a representational similarity matrix (RSM) in which a value of 1 was assigned to pairs of items of the same category and a value of zero to items of different categories (‘Category model’, Figure 2A). This model RSM was correlated with a time series of neural RSMs in each of our ROIs. Pairwise correlations among stimuli were computed in windows of 500ms, overlapping by 400ms, using the representational patterns described in the section above, resulting in an RSA time-frequency map in each of our ROIs. In order to obtain a robust estimate of the multivariate patterns representing individual items, we averaged the time-frequency activity across repetitions of items throughout the experiment in each channel independently before building the neural RSMs. RSM time-series were vectorized by removing the diagonal values and taking only half of the matrix given its symmetry at each time-frequency point. We correlated vectorized model RSMs and neural RSMs at each time-frequency point using Spearman’s rho, and evaluated whether the resulting isher Z-transformed rho-values were different from zero at the group level to determine statistical significance (two-sided tests). Multiple comparisons correction was performed using cluster-based permutation statistics (see below), and we Bonferroni corrected the final results to account for the number of layers tested in each network.

### Contrast-based RSA

In order to test the reoccurrence and stability of activity patterns in our two regions of interest during encoding and between encoding and maintenance, we performed two contrast-based pattern similarity analyses, as a complementary analysis to the model-based RSA approach (Figure 2). In particular, we investigated the presence of category-specific information in our data by contrasting correlations between different items of the same category with correlations between different items from different categories. This was done separately for items presented in different trials during encoding (encoding-encoding similarity, EES) and between encoding and maintenance (encoding-maintenance similarity, EMS). Only items belonging to different trials were included in this analysis to avoid any spurious correlations driven by the autocorrelation of the signal. Similar to the model-based RSA approach, we averaged across item repetitions before conducting the similarity comparisons.

We computed similarities for same-category and different-category item pairs and averaged across all combinations of items in the same condition in each subject independently (rho values were Fisher z-transformed before averaging). The same-category and different-category correlations were then statistically compared at the group level using *t*-tests. In the different-category condition, we excluded item pairs containing stimuli presented in overlapping trials after averaging, again to avoid any possible bias related to the autocorrelation of the signal. As an example, if a Robot exemplar was presented in trials 2, 4 and 8, and a planet was presented in trials 7, 9 and 8, the average correlation corresponding to these items would contain activity of an overlapping trial (8 in the example). The correlation corresponding to these two items would therefore not be included. Note that this was not necessary for the same category correlations. We quantified the similarity of neural representations by comparing epochs of brain activity separately in VVS and PFC. Note that contrary to the model-based RSA approach (see above) this analysis was not performed at each individual frequency but frequencies were grouped into five frequency bands. This effectively increased the information content (variance) of our representational patterns, making them more suitable to investigate their reoccurrence during encoding and maintenance. Moreover, combining individual frequencies into bands allowed us to reduce the dimensionality of the results when comparing all pairwise combinations of time points in the temporal generalization analysis. We computed the correlation of these representational patterns across all available time-points using a sliding time window approach proceeding in time steps of 100ms (i.e., with an 80% overlap). This resulted in a temporal generalization matrix with two temporal dimensions on the vertical and horizontal axes (Figure 2D and 2F). Note that values in these matrices reflect both lagged (off-diagonal) and non-lagged (on-diagonal) correlations and were thus informative about the stability of neural representations over time.

Pattern similarity maps were computed for each pair of items in a correspondent condition at each time-window and rho values in these maps were Fisher z-transformed for statistical analysis. The temporal generalization maps were averaged across conditions for each subject independently, and the resulting average maps were contrasted via paired *t*-tests across conditions at the group level.

Please note that in all pattern similarity plots (and also in the DNN-RSA plots, see below), correlations corresponding to each 500ms window were assigned to the time point at the center of the respective window (e.g., a time bin corresponding to activity from 0 to 500ms was assigned to 250ms).

### RSA at high temporal resolution

We increased the temporal resolution of our sliding time window approach to compare the onset of category-specific information in VVS and PFC (Figure 2E). In this analysis, power values were computed with the same method and parameters as in the main contrast-based analysis, but at an increased temporal resolution (10ms). Feature vectors were constructed in 500ms time windows and the 50 time-points included in each window were averaged separately for electrodes and frequencies, resulting in a two-dimensional representational pattern. We included all individual frequencies in the 3-150Hz range (a total of 52). These two-dimensional frequency x electrode vectors were concatenated into one-dimensional arrays for similarity analyses. We employed a sliding time-window approach with incremental steps of 10ms resulting in an overlap of 490ms between two consecutive windows, focusing only on matching time points (non-lagged correlations). We performed this analysis separately for the VVS and the PFC and assessed the statistical significance of the resulting time-series in each region. At each time point, we compared the group-level Fisher z-transformed rho values against zero. We also directly compared the values between PFC and VVS at the group level. Given that not all subjects had implanted electrodes in both of our two ROIs, we performed unpaired *t*-tests at each time-point. We corrected for multiple comparisons by applying cluster-based permutation statistics in the temporal dimension in all the pattern similarity analyses (see section *Multiple Comparisons Correction* below).

### Feedforward and recurrent DNN models

We compared VWM representations in the iEEG data with those formed in two types of convolutional deep neural network (DNN) architectures: feedforward and recurrent DNNs. We used AlexNet (Krizhevsky et al., 2012), a widely applied network in computational cognitive neuroscience to model visual perception and WM, as our feedforward model (Kuzovkin et al., 2018; Liu et al., 2020). We also employed two recurrent convolutional DNNs: BL-NET, which has been recently applied to model human reaction times in a perceptual recognition task (Spoerer et al., 2020), and corNET-RT, a network recently developed to model information processing in the primate ventral visual stream (Kubilius et al., 2018). AlexNet is a deep convolutional feedforward neural network composed of five convolutional layers and 3 fully connected layers that simulates the hierarchical structure of neurons along the ventral visual stream. AlexNet was trained in the task of object identification, i.e., the assignment of object labels to visual stimuli, using the ImageNet dataset (Deng et al., 2009). When learning to identify images, AlexNet develops layered representations of stimuli that hierarchically encode increasingly abstract visual properties: Early layers reflect low-level features of images such as edges or textures while deeper layers are sensitive to more complex visual information, such as the presence of objects or object parts. Several studies demonstrated the validity of AlexNet as a model of neural representations during biological vision, showing that it can capture relevant features of information processing in the VVS of humans during perceptual and mnemonic processing (Bao et al., 2020; Kuzovkin et al., 2018; Liu et al., 2021). We computed RSMs at every convolutional and fully connected layers of the network, following previous work (Liu et al., 2020, 2021).

The BL-NET is a deep recurrent convolutional neural network consisting of 7 convolutional layers with feedforward and lateral recurrent connections, followed by 7 batch normalization and RELU layers. Every unit in the BL-NET network receives lateral input from other units within feature maps. BL-NET has demonstrated high accuracy in the task of object recognition (Spoerer et al., 2020) after being trained with two large-scale image datasets (i.e., ImageNet and Ecoset; Mehrer et al., 2021; Spoerer et al., 2020). We tested the network trained with these two different datasets in our analyses. Given that the output of each layer, which combines activity of lateral and feedforward connections, is computed at every single time-step in the RELU layers of the model, we selected these specific layers to compute the RSMs in our main analyses (Spoerer et al., 2020). We obtained similar results when we compared the activations extracted from the convolutional layers (after batch normalization).

The corNET-RT network is another prominent example of recurrent architectures that have been employed to model neural activity in the VVS of primates. It comprises four layers designed to capture information processing in the main four VVS regions: V1, V2, V4, and IT. Like the BL-NET, corNET-RT exclusively incorporates lateral and not across area connectivity. Each layer of the network consists of an input and output convolutional layer, group normalization and RELU non-linearities. Unlike the BL-NET, the number of recurrent steps in each layer is not fixed but varies from 5 (in layer V1) to 2 (in layer IT). RSMs were computed specifically for the convolutional layers (we selected the output convolutional modules in each layer), although similar results were observed when RSMs were computed from the outputs of the non-linear layers.

BL-NET and corNET-RT are two of the most prominent task-performing convolutional DNN models for image classification that have introduced recurrence as a main architectural feature. These networks have shown improvements in performance as compared to parameter-matched feedforward networks in the complex task of object recognition (Kubilius et al., 2018; Spoerer et al., 2020). Theoretical accounts and experimental findings have proposed that recurrent DNNs can better explain neural activity in the VVS and behavioral data than feedforward networks (Kar et al., 2019; Kietzmann et al., 2019; Kubilius et al., 2018; Nayebi et al., 2018; H. Tang et al., 2018). While previous studies characterized neural representations in humans using recurrent models in the domain of visual perception (Kietzmann et al., 2019), no study so far has used these types of architectures to model VWM, and no study has applied them to iEEG data.

Note that the BL-NET and the corNET-RT networks have different unrolling schemes across time, which affects how activity propagates through the networks. In BL-NET, feedforward and recurrent processing happen in parallel: a feedforward pass takes no time, while each recurrent step takes 1 time point. Thus, each layer receives a time-varying feedforward input. In corNET-RT, on the other hand, the onset of responses at deep layers is delayed when recurrence is engaged in earlier layers. These two approaches have been referred to as unrolling in ‘biological’ time (corNET-RT) vs ‘engineering’ time (BL-NET; Kubilius et al., 2018; Spoerer et al., 2020).

### Stimulus representations in DNNs

In order to analyze how the different DNN architectures represented the stimuli in our study, we presented the networks with our images and computed unit activations at each layer. We calculated Spearman’s correlations between the D features for every pair of pictures, resulting in a 60 × 60 representational similarity matrix (RSM) in each layer (Kriegeskorte et al., 2008). The Alexnet unit activations were computed using the Matlab Deep Learning Toolbox. Images were scaled to fit the 227×227 input layer of the network. The unit activations in the BL-NET network were extracted using the pipeline described in https://github.com/cjspoerer/rcnn-sat. Images were scaled to 128×128 pixels, and normalized to values between -1 and 1 to fit the input layer of the network as it was originally trained. The number of recurrent passes in the BL-NET architecture was set to 8 time-steps in each layer. We extracted the unit activations at each of these time points and computed RSMs, resulting in a total of 7 (layers) x 8 (time-points) = 56 RSMs. corNET-RT activations were extracted using TorchLens (Taylor & Kriegeskorte, 2023), and we corroborated the results using the ‘Torch ision’ toolbox (Muttenthaler & Hebart, 2021). Images were z-scored to the mean and standard deviation of the ImageNet database and scaled to 224×224 pixels to match the training parameters of the network.

To visualize the representations of stimuli in our networks, we employed multi-dimensional scaling (MDS). MDS is a dimensionality reduction technique which exploits the geometric properties of RSMs, projecting the high-dimensional network activation patterns into lower-dimensional spaces. To apply the MDS algorithm to our RSMs, we subtracted the correspondent values in the matrix from 1 to obtain a distance metric and projected the data into two dimensions (Figures 3A, 4D and 5A).

Importantly, all three architectures we employed were trained with the Imagenet dataset, in which none of the categories included in our study (‘house’, ‘robot’, ‘hand’, ‘face’, ‘planet’, and tree’) are present as object labels. For this reason, we did not focus our analysis on network classification performance but characterized categorical representations that were formed across layers, computing within-category, between-category correlations and their difference (CCI scores, see below). Moreover, we performed an additional analysis involving a variant of BL-NET trained with the Ecoset dataset (Mehrer et al., 2021), which contains part of our stimuli labels (i.e., labels ‘house’, ‘robot’ and ‘tree’) to corroborate our main results. Note that the Ecoset analysis was specifically performed in the clusters where we observed significant time-frequency fits in the main analysis: all layers during encoding and layers 4, 5 and 6 during maintenance in the VVS; only layer 7 during maintenance in the PFC. For each subject, we computed the average correlation in the time-frequency cluster identified in the main analysis and contrasted these average rho values against zero at the group level to determine statistical significance (Supplementary figure 2).

In addition to computing the representational geometry of stimuli across all layers of the networks, we uantified the similarity between RSMs across layers using Spearman’s Rho (a “second-level” similarity metric; Mehrer et al., 2020). We applied MDS to visualize the similarity structure of the initial and last time points in each layer of BL-NET and Cornet-RT (Figure 4B and 5B).

To quantify the amount of category information in the different layers of the networks, we computed a Category Cluster Index (CCI), defined as the difference of average within-category and between-category correlations in the DNN representations of the stimuli. Both within and across category correlation averages were computed after removing the diagonal of the RSM matrices (which only contains values of 1 by definition) and duplicated values due to the symmetry of the RSMs. CCI approaches 1 if representations in all categories are perfectly clustered and 0 if no categorical structure is present in the data (Mehrer et al., 2020). We computed CCI at each layer of the Alexnet (Figure 3C), and for each time point in each layer of the BL-Net and the corNET-RT networks (Figure 4D and 5D). To assess whether the observed CCI values were significant, we implemented a permutation procedure. We built a distribution of CCI values expected by chance by shuffling the trial labels of the network RSMs 1,000 times and recomputing CCI values. We considered significant CCI values that exceeded the 95^th^ percentile of these null distributions.

In order to better characterize categorical representations in our networks and directly compare them, we performed a linear fit of within-category and between-category correlations across layers (Supplementary Figure 1). We specifically focused on the last time point in each layer in our recurrent architectures. We computed the correlation of the activations corresponding to every pair of items in each layer and performed a linear least-squares fit with the resulting values (270 within-category correlations and 1,500 between-category correlations were computed in each layer). To evaluate whether correlations increased or decreased linearly across layers, we compared the distribution of slopes taken from the linear fit against zero (representing the null hypothesis of an average flat line) in each individual network. In addition, we compared these distributions across networks using paired t-tests.

Please note that given that two versions of the experiment were created for German and Chinese participants (with Angela Merkel and Jackie Chan as face stimuli, respectively), we passed through the networks two different datasets of images. For visualization of network RSMs and corresponding MDS plots (Figures 3A, 4E and 5E), we employed the German version of the stimuli. In the analyses focusing on the network representations, we generated independent statistics for the two stimuli sets and then averaged them. This applies to the plots showing the representational consistency of networks across layers and time-points, within- and between-category correlations and CCI scores (Figures 3B-C, 4A-D, and 5A-D).

### Representational similarity analyses based on DNNs: Modeling neural representations with deep neural networks

We compared the representations formed in the DNN architectures with the iEEG representations using RSA. Neural RSMs were constructed following the procedure described in the section *Model-based RSA* above.

Similar to the category-model analysis, we performed a time-frequency resolved analysis of fits of neural and DNN-based RSMs. In this analysis, neural RSM time series (same time windows as described above) were computed with feature vectors comprising information of each individual fre uency independently (e.g., for 3Hz, 4Hz, … 150Hz). The resulting RSM time-series were correlated with network RSMs at each individual layer. Individual frequencies were extracted using the same parameters as in the category-model and the contrast-based analyses (see section *Time-frequency analysis*). The resulting time-series of correlation values were stacked into time-frequency maps of model fits (Figure 4I and 5I). Significance was determined by contrasting the observed Fischer-Z transformed rho values against zero at the group level. Results were corrected for multiple comparisons using cluster-based permutation statistics and we additionally applied Bonferroni corrections across layers (see below).

### Multiple comparisons corrections

We performed cluster-based permutation statistics to correct for multiple comparisons in the pattern similarity analyses (Figure 2) and in the RSA-DNN analyses (Figures 3-5).

In the pattern similarity analyses, we applied cluster-based permutation statistics both for the temporal generalization analysis (Figure 2D and 2F), and for the temporally resolved analysis (Figure 2E). For both analyses, we contrasted same and different category correlations at different time-points using *t*-tests, as in the main analysis, after shuffling the trial labels 1,000 times. We considered significant a time-point if the difference between these surrogate conditions was significant at *p* < 0.05 (two-tailed tests were employed). At every permutation, we computed clusters of significant values defined as contiguous regions in time where significant correlations were observed and took the largest cluster at each permutation. Please note that in the temporal generalization analysis, time was defined in two dimensions and clusters were formed by grouping significant values across both of these dimensions, while in the temporally resolved analysis (Figure 2E), correlations were computed at matching time-points and clusters were formed along one temporal dimension. In both analyses, the permutation procedure resulted in a distribution of surrogate *t*-values under the assumption of the null hypothesis. We only considered significant those contiguous time pairs in the empirical (non-shuffled) data whose summed *t*-values exceeded the summed *t*-value of 95% of the distribution of surrogate clusters (corresponding to a corrected P < 0.05; Maris & Oostenveld, 2007).

We also performed cluster-based permutation statistics in the analysis at high temporal resolution in which we directly compared similarity values between VVS and PFC. In this analysis, we computed clusters of significant EES differences between the two regions for every time-point by applying unpaired *t*-tests. We repeated this analysis 1,000 times after shuffling the region labels and kept the summed value of the largest cluster at every permutation. We only considered significant those clusters in the empirical data above the 95^th^ percentile of the shuffled distribution. In the RSA-DNN analyses (and also in the category-model RSA analysis, Figure 2), we applied cluster-based permutation statistics. To determine the significance of the correlations between neural and model RSMs, we recalculated the model RSMs at each layer of the network after randomly shuffling the labels of the images. The surrogate model similarity matrices were then correlated with the neural similarity matrix 1,000 times at all time-frequency pairs. As in the original analysis, we computed the correlations after removing the diagonal of the RSMs and only took half of the matrices given their symmetry. We identified clusters of contiguous windows in the time-frequency domain where the group-level correlations between neural and network RSMs were significantly different from zero at p < 0.05 (two-sided test) and selected the maximum cluster size of summed *t*-values for every permutation. This resulted in a distribution of surrogate *t*-values. The statistical significance was then determined by comparing the correlation values for the empirical data with the distribution of correlation values for the surrogate data (clusters whose summed *t*-values exceeded the 95% of the null distribution were considered significant).

In addition to cluster-based permutations, we also corrected our results for multiple comparisons using the Bonferroni method in the contrast-based RSA analyses (Figure 2) and in the model-based RSA analyses (Figures 2, 3, 4, and 5). In the contrast-based analyses, given that we tested five different frequency bands, we only considered *p*-values significant that were below an alpha of 0.05/5. In the model-based analyses, we adjusted the significance threshold according to the correspondent number of layers in each network that was tested (AlexNet= 8; BL-NET = 7, corNET-RT = 4). The same correction by number of layers was applied in the CCI analysis (Figure 3C, 4D and 5D).

### Data Availability

Anonymized intracranial EEG data and custom-written Matlab and Python code supporting the findings of this study will be made available upon publication.

## Acknowledgments

We would like to acknowledge DFG funding via the ORA project “WMREPS Hidden brain states underlying efficient representations in working memory” (project number 3 6 4 56). This project was a collaboration with Mark Stokes (Oxford) and Elkan Akyürek (Groningen) and is dedicated to the memory of Prof. Stokes.

## References

Antzoulatos, E. G., & Miller, E. K. (2014). Increases in functional connectivity between prefrontal cortex and striatum during category learning. Neuron, 83(1), 216–225.

Antzoulatos, E. G., & Miller, E. K. (2016). Synchronous beta rhythms of frontoparietal networks support only behaviorally relevant representations. Elife, 5, e17822.

Axmacher, N. (2020). Representational formats in medial temporal lobe and neocortex also determine subjective memory features. The Behavioral and Brain Sciences, 42, e283. 10.1017/S0140525X19001882

Baek, S., Song, M., Jang, J., Kim, G., & Paik, S.-B. (2021). Face detection in untrained deep neural networks. Nature Communications, 12(1), 7328. 10.1038/s41467-021-27606-9

Bao, P., She, L., McGill, M., & Tsao, D. Y. (2020). A map of object space in primate inferotemporal cortex. Nature, 583(7814), 103–108. 10.1038/s41586-020-2350-5

Barak, O., & Tsodyks, M. (2014). Working models of working memory. Current Opinion in Neurobiology, 25, 20–24. 10.1016/j.conb.2013.10.008

Bouchacourt, F., & Buschman, T. J. (2019). A flexible model of working memory. Neuron, 103(1), 147– 160.

Breedlove, J. L., St-Yves, G., Olman, C. A., & Naselaris, T. (2020). Generative Feedback Explains Distinct Brain Activity Codes for Seen and Mental Images. Current Biology, 30(12), 2211–2224.e6. 10.1016/j.cub.2020.04.014

Buschman, T. J., Denovellis, E. L., Diogo, C., Bullock, D., & Miller, E. K. (2012). Synchronous oscillatory neural ensembles for rules in the prefrontal cortex. Neuron, 76(4), 838–846.

Buschman, T. J., & Kastner, S. (2015). From behavior to neural dynamics: An integrated theory of attention. Neuron, 88(1), 127–144.

Buschman, T. J., & Miller, E. K. (2007). Top-down versus bottom-up control of attention in the prefrontal and posterior parietal cortices. Science, 315(5820), 1860–1862.

Cadieu, C. F., Hong, H., Yamins, D. L., Pinto, N., Ardila, D., Solomon, E. A., Majaj, N. J., & DiCarlo, J. J. (2014). Deep neural networks rival the representation of primate IT cortex for core visual object recognition. PLoS Computational Biology, 10(12), e1003963.

Chatham, C. H., Frank, M. J., & Badre, D. (2014). Corticostriatal output gating during selection from working memory. Neuron, 81(4), 930–942.

Cichy, R. M., Khosla, A., Pantazis, D., Torralba, A., & Oliva, A. (2016). Comparison of deep neural networks to spatio-temporal cortical dynamics of human visual object recognition reveals hierarchical correspondence. Scientific Reports, 6(1), 27755. 10.1038/srep27755

Cichy, R. M., Pantazis, D., & Oliva, A. (2014). Resolving human object recognition in space and time. Nature Neuroscience, 17(3), 455–462. 10.1038/nn.3635

Compte, A., Brunel, N., Goldman-Rakic, P. S., & Wang, X.-J. (2000). Synaptic Mechanisms and Network Dynamics Underlying Spatial Working Memory in a Cortical Network Model. Cerebral Cortex, 10(9), 910–923. 10.1093/cercor/10.9.910

Conwell, C., Jacob S. Prince, Kendrick N. Kay, George A. Alvarez, & Talia Konkle. (2023). What can 1.8 billion regressions tell us about the pressures shaping high-level visual representation in brains and machines? bioRxiv, 2022.03.28.485868. 10.1101/2022.03.28.485868

Delorme, A., & Makeig, S. (2004). EEGLAB: an open source toolbox for analysis of single-trial EEG dynamics including independent component analysis. Journal of Neuroscience Methods, 134(1), 9–21. 10.1016/j.jneumeth.2003.10.009

Deng, J., Dong, W., Socher, R., Li, L.-J., Li, K., & Fei-Fei, L. (2009). Imagenet: A large-scale hierarchical image database. 248–255.

D’Esposito, M. (2007). From cognitive to neural models of working memory. Philosophical Transactions of the Royal Society B: Biological Sciences, 362(1481), 761–772. 10.1098/rstb.2007.2086

D’Esposito, M., Detro, J. A., Alsop, D. C., Shin, R. K., Atlas, S., & Grossman, M. (1995). The neural basis of the central executive system of working memory. Nature, 378(6554), 279–281. 10.1038/378279a0

Ehrlich, D. B., & Murray, J. D. (2022). Geometry of neural computation unifies working memory and planning. Proceedings of the National Academy of Sciences, 119(37), e2115610119. 10.1073/pnas.2115610119

Eichenbaum, H. (2017). Memory: Organization and Control. Annual Review of Psychology, 68, 19–45. 10.1146/annurev-psych-010416-044131

Engel, A. K., & Fries, P. (2010). Beta-band oscillations—Signalling the status quo? Current Opinion in Neurobiology, 20(2), 156–165.

Ester, E. F., Nouri, A., & Rodriguez, L. (2018). Retrospective cues mitigate information loss in human cortex during working memory storage. Journal of Neuroscience, 38(40), 8538–8548.

Fellner, M.-C., Waldhauser, G. T., & Axmacher, N. (2020). Tracking selective rehearsal and active inhibition of memory traces in directed forgetting. Current Biology, 30(13), 2638–2644.

Gelastopoulos, A., Whittington, M. A., & Kopell, N. J. (2019). Parietal low beta rhythm provides a dynamical substrate for a working memory buffer. Proceedings of the National Academy of Sciences, 116(33), 16613–16620. 10.1073/pnas.1902305116

Griffin, I. C., & Nobre, A. C. (2003). Orienting attention to locations in internal representations. Journal of Cognitive Neuroscience, 15(8), 1176–1194.

Heinen, R., Bierbrauer, A., Wolf, O. T., & Axmacher, N. (2023). Representational formats of human memory traces. Brain Structure & Function. 10.1007/s00429-023-02636-9

Higo, T., Mars, R. B., Boorman, E. D., Buch, E. R., & Rushworth, M. F. S. (2011). Distributed and causal influence of frontal operculum in task control. Proceedings of the National Academy of Sciences, 108(10), 4230–4235. 10.1073/pnas.1013361108

Kar, K., & DiCarlo, J. J. (2021). Fast recurrent processing via ventrolateral prefrontal cortex is needed by the primate ventral stream for robust core visual object recognition. Neuron, 109(1), 164–176.

Kar, K., Kubilius, J., Schmidt, K., Issa, E. B., & DiCarlo, J. J. (2019). Evidence that recurrent circuits are v m’ u f bj b v Nature Neuroscience, 22(6), 974–983.

Kerren, C., Linde-Domingo, J., & Spitzer, B. (2022). Prioritization of semantic over visuo-perceptual aspects in multi-item working memory. bioRxiv.

Khaligh-Razavi, S.-M., & Kriegeskorte, N. (2014). Deep supervised, but not unsupervised, models may explain IT cortical representation. PLoS Computational Biology, 10(11), e1003915.

Kietzmann, T. C., Spoerer, C. J., Sörensen, L. K., Cichy, R. M., Hauk, O., & Kriegeskorte, N. (2019). Recurrence is required to capture the representational dynamics of the human visual system. Proceedings of the National Academy of Sciences, 116(43), 21854–21863.

Kriegeskorte, N., & Diedrichsen, J. (2019). Peeling the Onion of Brain Representations. Annual Review of Neuroscience, 42, 407–432. 10.1146/annurev-neuro-080317-061906

Kriegeskorte, N., Mur, M., & Bandettini, P. A. (2008). Representational similarity analysis-connecting the branches of systems neuroscience. Frontiers in Systems Neuroscience, 4.

Krizhevsky, A., Sutskever, I., & Hinton, G. E. (2012). Imagenet classification with deep convolutional neural networks. Communications of the ACM, 60(6), 84–90.

Kubilius, J., Schrimpf, M., Nayebi, A., Bear, D., Yamins, D. L., & DiCarlo, J. J. (2018). Cornet: Modeling the neural mechanisms of core object recognition. BioRxiv, 408385.

Kuzovkin, I., Vicente, R., Petton, M., Lachaux, J.-P., Baciu, M., Kahane, P., Rheims, S., Vidal, J. R., & Aru, J. (2018). Activations of deep convolutional neural networks are aligned with gamma band activity of human visual cortex. Communications Biology, 1(1), 1–12.

Kwak, Y., & Curtis, C. E. (2022). Unveiling the abstract format of mnemonic representations. Neuron.

Lepsien, J., & Nobre, A. C. (2006). Cognitive control of attention in the human brain: Insights from orienting attention to mental representations. Brain Research, 1105(1), 20–31.

Lepsien, J., Thornton, I., & Nobre, A. C. (2011). Modulation of working-memory maintenance by directed attention. Neuropsychologia, 49(6), 1569–1577.

Liebe, S., Hoerzer, G. M., Logothetis, N. K., & Rainer, G. (2012). Theta coupling between V4 and prefrontal cortex predicts visual short-term memory performance. Nature Neuroscience, 15(3), 456–462. 10.1038/nn.3038

Lindsay, G. W. (2021). Convolutional neural networks as a model of the visual system: Past, present, and future. Journal of Cognitive Neuroscience, 33(10), 2017–2031.

Liu, J., Zhang, H., Yu, T., Ni, D., Ren, L., Yang, Q., Lu, B., Wang, D., Heinen, R., Axmacher, N., & others. (2020). Stable maintenance of multiple representational formats in human visual short-term memory. Proceedings of the National Academy of Sciences, 117(51), 32329–32339.

Liu, J., Zhang, H., Yu, T., Ren, L., Ni, D., Yang, Q., Lu, B., Zhang, L., Axmacher, N., & Xue, G. (2021). Transformative neural representations support long-term episodic memory. Science Advances, 7(41), eabg9715.

Lundqvist, M., Herman, P., Warden, M. R., Brincat, S. L., & Miller, E. K. (2018). Gamma and beta bursts during working memory readout suggest roles in its volitional control. Nature Communications, 9(1), 1–12.

Mante, V., Sussillo, D., Shenoy, K. V., & Newsome, W. T. (2013). Context-dependent computation by recurrent dynamics in prefrontal cortex. Nature, 503(7474), 78–84. 10.1038/nature12742

Maris, E., & Oostenveld, R. (2007). Nonparametric statistical testing of EEG-and MEG-data. Journal of Neuroscience Methods, 164(1), 177–190.

McKee, J. L., Riesenhuber, M., Miller, E. K., & Freedman, D. J. (2014). Task dependence of visual and category representations in prefrontal and inferior temporal cortices. Journal of Neuroscience, 34(48), 16065–16075.

Mehrer, J., Spoerer, C. J., Jones, E. C., Kriegeskorte, N., & Kietzmann, T. C. (2021). An ecologically motivated image dataset for deep learning yields better models of human vision. Proceedings of the National Academy of Sciences, 118(8), e2011417118. 10.1073/pnas.2011417118

Mehrer, J., Spoerer, C. J., Kriegeskorte, N., & Kietzmann, T. C. (2020). Individual differences among deep neural network models. Nature Communications, 11(1), 1–12.

Miller, E. K., & Cohen, J. D. (2001). An integrative theory of prefrontal cortex function. Annual Review of Neuroscience, 24, 167–202. 10.1146/annurev.neuro.24.1.167

Miller, E. K., Lundqvist, M., & Bastos, A. M. (2018). Working Memory 2.0. Neuron, 100(2), 463–475.

Muttenthaler, L., & Hebart, M. N. (2021). THINGSvision: A Python Toolbox for Streamlining the Extraction of Activations From Deep Neural Networks. Frontiers in Neuroinformatics, 15. https://www.frontiersin.org/articles/10.3389/fninf.2021.679838

Myers, N. E., Stokes, M. G., & Nobre, A. C. (2017). Prioritizing information during working memory: Beyond sustained internal attention. Trends in Cognitive Sciences, 21(6), 449–461.

Nayebi, A., Bear, D., Kubilius, J., Kar, K., Ganguli, S., Sussillo, D., DiCarlo, J. J., & Yamins, D. L. (2018). Task-driven convolutional recurrent models of the visual system. Advances in Neural Information Processing Systems, 31.

Nee, D. E., & Jonides, J. (2008). Neural correlates of access to short-term memory. Proceedings of the National Academy of Sciences, 105(37), 14228–14233. 10.1073/pnas.0802081105

Nee, D. E., & Jonides, J. (2009). Common and distinct neural correlates of perceptual and memorial selection. Neuroimage, 45(3), 963–975.

Nelissen, N., Stokes, M., Nobre, A. C., & Rushworth, M. F. S. (2013). Frontal and parietal cortical interactions with distributed visual representations during selective attention and action selection. Journal of Neuroscience, 33(42), 16443–16458.

Nobre, A. C., Coull, J. T., Maquet, P., Frith, C. D., Vandenberghe, R., & Mesulam, M. M. (2004). Orienting attention to locations in perceptual versus mental representations. Journal of Cognitive Neuroscience, 16(3), 363–373.

Oostenveld, R., Fries, P., Maris, E., & Schoffelen, J.-M. (2011). FieldTrip: Open source software for advanced analysis of MEG, EEG, and invasive electrophysiological data. Computational Intelligence and Neuroscience, 2011, 156869. 10.1155/2011/156869

Pacheco Estefan, D., Sánchez-Fibla, M., Duff, A., Principe, A., Rocamora, R., Zhang, H., Axmacher, N., & Verschure, P. F. M. J. (2019). Coordinated representational reinstatement in the human hippocampus and lateral temporal cortex during episodic memory retrieval. Nature Communications, 10(1), 1–13.

Pacheco Estefan, D., Zucca, R., Arsiwalla, X., Principe, A., Zhang, H., Rocamora, R., Axmacher, N., & Verschure, P. F. M. J. (2021). Volitional learning promotes theta phase coding in the human hippocampus. Proceedings of the National Academy of Sciences, 118(10).

Panichello, M. F., & Buschman, T. J. (2021). Shared mechanisms underlie the control of working memory and attention. Nature, 592(7855), 601–605.

Posner, M. I. (1980). Orienting of attention. Quarterly Journal of Experimental Psychology, 32(1), 3–25.

Rissman, J., & Wagner, A. D. (2012). Distributed representations in memory: Insights from functional brain imaging. Annual Review of Psychology, 63, 101–128. 10.1146/annurev-psych-120710-100344

Schmidt, B. K., Vogel, E. K., Woodman, G. F., & Luck, S. J. (2002). Voluntary and automatic attentional control of visual working memory. Perception & Psychophysics, 64(5), 754–763.

Sörensen, L. K. A., Bohté, S. M., de Jong, D., Slagter, H. A., & Scholte, H. S. (2023). Mechanisms of human dynamic object recognition revealed by sequential deep neural networks. PLOS Computational Biology, 19(6), e1011169. 10.1371/journal.pcbi.1011169

Spitzer, B., & Blankenburg, F. (2011). Stimulus-dependent EEG activity reflects internal updating of tactile working memory in humans. Proceedings of the National Academy of Sciences, 108(20), 8444–8449. 10.1073/pnas.1104189108

Spitzer, B., Evelin Wacker, & Felix Blankenburg. (2010). Oscillatory Correlates of Vibrotactile Frequency Processing in Human Working Memory. The Journal of Neuroscience, 30(12), 4496. 10.1523/JNEUROSCI.6041-09.2010

Spitzer, B., Fleck, S., & Blankenburg, F. (2014). Parametric alpha-and beta-band signatures of supramodal numerosity information in human working memory. Journal of Neuroscience, 34(12), 4293– 4302.

Spitzer, B., & Haegens, S. (2017). Beyond the Status Quo: A Role for Beta Oscillations in Endogenous Content (Re)Activation. Eneuro, 4(4), ENEURO.0170-17.2017. 10.1523/ENEURO.0170-17.2017

Spoerer, C. J., Kietzmann, T. C., Mehrer, J., Charest, I., & Kriegeskorte, N. (2020). Recurrent neural networks can explain flexible trading of speed and accuracy in biological vision. PLoS Computational Biology, 16(10), e1008215.

Sprague, T. C., Ester, E. F., & Serences, J. T. (2016). Restoring latent visual working memory representations in human cortex. Neuron, 91(3), 694–707.

Stanley, D. A., Roy, J. E., Aoi, M. C., Kopell, N. J., & Miller, E. K. (2018). Low-beta oscillations turn up the gain during category judgments. Cerebral Cortex, 28(1), 116–130.

Staresina, B. P., Michelmann, S., Bonnefond, M., Jensen, O., Axmacher, N., & Fell, J. (2016). Hippocampal pattern completion is linked to gamma power increases and alpha power decreases during recollection. eLife, 5, e17397. 10.7554/eLife.17397

Stokes, M. G., Kusunoki, M., Sigala, N., Nili, H., Gaffan, D., & Duncan, J. (2013). Dynamic coding for cognitive control in prefrontal cortex. Neuron, 78(2), 364–375.

Stolk, A., Griffin, S., van der Meij, R., Dewar, C., Saez, I., Lin, J. J., Piantoni, G., Schoffelen, J.-M., Knight, R. T., & Oostenveld, R. (2018). Integrated analysis of anatomical and electrophysiological human intracranial data. Nature Protocols, 13(7), 1699–1723. 10.1038/s41596-018-0009-6

Tang, H., Schrimpf, M., Lotter, W., Moerman, C., Paredes, A., Ortega Caro, J., Hardesty, W., Cox, D., & Kreiman, G. (2018). Recurrent computations for visual pattern completion. Proceedings of the National Academy of Sciences, 115(35), 8835–8840. 10.1073/pnas.1719397115

Tang, W., Shin, J. D., & Jadhav, S. P. (2023). Geometric transformation of cognitive maps for generalization across hippocampal-prefrontal circuits. Cell Reports, 42(3), 112246. 10.1016/j.celrep.2023.112246

Taylor, J., & Kriegeskorte, N. (2023). TorchLens: A Python package for extracting and visualizing hidden activations of PyTorch models. bioRxiv, 2023.03.16.532916. 10.1101/2023.03.16.532916

Ten Oever, S., Sack, A. T., Oehrn, C. R., & Axmacher, N. (2021). An engram of intentionally forgotten information. Nature Communications, 12(1), 6443. 10.1038/s41467-021-26713-x

van Bergen, R. S., & Kriegeskorte, N. (2020). Going in circles is the way forward: The role of recurrence in visual inference. Current Opinion in Neurobiology, 65, 176–193.

Vezoli, J., Vinck, M., Bosman, C. A., Bastos, A. M., Lewis, C. M., Kennedy, H., & Fries, P. (2021). Brain rhythms define distinct interaction networks with differential dependence on anatomy. Neuron, 109(23), 3862–3878.

Vinken, K., & Op de Beeck, H. (2021). Using deep neural networks to evaluate object vision tasks in rats. PLOS Computational Biology, 17(3), e1008714. 10.1371/journal.pcbi.1008714

Vogel, E. K., & Machizawa, M. G. (2004). Neural activity predicts individual differences in visual working memory capacity. Nature, 428(6984), 748–751. 10.1038/nature02447

Wallis, G., Stokes, M., Cousijn, H., Woolrich, M., & Nobre, A. C. (2015). Frontoparietal and cingulo-opercular networks play dissociable roles in control of working memory. Journal of Cognitive Neuroscience, 27(10), 2019–2034.

Wan, Q., Menendez, J. A., & Postle, B. R. (2022). Priority-based transformations of stimulus representation in visual working memory. PLoS Computational Biology, 18(6), e1009062. 10.1371/journal.pcbi.1009062

Wang, X. J. (2001). Synaptic reverberation underlying mnemonic persistent activity. Trends in Neurosciences, 24(8), 455–463. 10.1016/s0166-2236(00)01868-3

Wang, X.-J. (2001). Synaptic reverberation underlying mnemonic persistent activity. Trends in Neurosciences, 24(8), 455–463.

Weber, J., Iwama, G., Solbakk, A.-K., Blenkmann, A. O., Larsson, P. G., Ivanovic, J., Knight, R. T., Endestad, T., & Helfrich, R. (2023). Subspace partitioning in the human prefrontal cortex resolves cognitive interference. Proceedings of the National Academy of Sciences, 120(28), e2220523120. 10.1073/pnas.2220523120

Wimmer, K., Marc Ramon, Tatiana Pasternak, & Albert Compte. (2016). Transitions between Multiband Oscillatory Patterns Characterize Memory-Guided Perceptual Decisions in Prefrontal Circuits. The Journal of Neuroscience, 36(2), 489. 10.1523/JNEUROSCI.3678-15.2016

Wu, X., & Fuentemilla, L. (2023). Distinct encoding and post-encoding representational formats contribute to episodic sequence memory formation. Cerebral Cortex (New York, N.Y. : 1991), bhad138. 10.1093/cercor/bhad138

Xue, G. (2022). From remembering to reconstruction: The transformative neural representation of episodic memory. Progress in Neurobiology, 219, 102351. 10.1016/j.pneurobio.2022.102351

Yamins, D. L., & DiCarlo, J. J. (2016). Using goal-driven deep learning models to understand sensory cortex. Nature Neuroscience, 19(3), 356–365.

Yamins, D. L., Hong, H., Cadieu, C. F., Solomon, E. A., Seibert, D., & DiCarlo, J. J. (2014). Performance-optimized hierarchical models predict neural responses in higher visual cortex. Proceedings of the National Academy of Sciences, 111(23), 8619–8624.

